# Reverse engineering lateral root stable prebranch site formation; Complementary roles for auxin and auxin signalling

**DOI:** 10.1101/2022.06.24.497450

**Authors:** Joana Teixeira Santos, Thea van den Berg, Kirsten ten Tusscher

**Affiliations:** Computational Developmental Biology Group, Faculty of Science, Utrecht University, The Netherlands

**Keywords:** lateral roots, prebranch sites, priming, auxin, auxin signalling, temporal integration

## Abstract

Priming is the process through which periodic elevations in auxin signalling prepattern future sites for lateral root formation, called prebranch sites. Thusfar is has remained a matter of debate to what extent elevations in auxin concentration and/or auxin signalling are critical for priming and prebranch site formation. Recently, we discovered a reflux-and-growth mechanism for priming generating periodic elevations in auxin concentration that subsequently dissipate. Here we reverse engineer a mechanism for prebranch site formation that translates these transient elevations into a persistent increase in auxin signalling, resolving the prior debate into a two-step process of auxin concentration mediated initial signal and auxin signalling capacity mediated memorization. A critical aspect of the prebranch site formation mechanism is its activation in response to time integrated rather than instantaneous auxin signalling. The proposed mechanism is demonstrated to be consistent with prebranch site auxin signalling dynamics, lateral inhibition and symmetry breaking mechanisms and perturbations in auxin homeostasis.

**Summary statement:** Using computational modeling we reveal the likely complementary roles of auxin and auxin signalling in one of the earliest step in the formation of plant lateral roots, prebranch site formation.

## Introduction

The architecture of the plant root system -length of the main root, number, length, positioning and angles of lateral roots - determines its access to water and nutrients. As a consequence, root system architecture (RSA) is a major determinant of plant fitness and crop yields (Herder et al. 2010; Rogers and Benfey 2015). Being the plants hidden half, its limited accessibility has caused the root system to have been far less subjected to targeted breeding efforts. Optimization of crop species root systems is therefore considered to be one of the most promising targets for achieving a next green revolution (Herder et al. 2010; Kong et al. 2014; Wollenweber, Porter, and Lübberstedt 2005). In order to achieve this, an in depth understanding of the developmental processes shaping root system architecture and the mechanisms enabling its plastic adjustment to environmental conditions is needed. Still, many aspects of RSA patterning remain to be revealed.

A critical aspect in RSA patterning is the formation of lateral roots. LR development starts with a process called priming, the prepatterning of sites competent for future lateral root formation. This process entails periodic oscillations in auxin levels and/or auxin signalling at the end of the meristem and start of the elongation zone (Fig 1A) (De Smet et al. 2007; Moreno-Risueno et al. 2010; Xuan et al. 2015, 2016), that through growth become translated into a spatially periodic pattern of lateral root competent sites (Moreno-Risueno et al. 2010; Xuan et al. 2015, 2016). Only if priming has sufficient amplitude it is successful and leads to the formation of a stable prebranch site (PBS) (Fig 1B,C) (Xuan et al. 2015), after which founder cell identity establishment and LR initiation may occur. After this, these cells asymmetrically divide, initiating the actual LR formation process. In the subsequent stage, the overlaying endodermal tissue layer is penetrated by the LRP and after this cortical and epidermal cells are pushed aside, so that the primordium emerges (Malamy and Benfey 1997). Before the LR starts to elongate, the meristem of the new LR becomes activated. LR will eventually recapitulate the pattern of developmental zones seen in the main root.

**Fig. 1.**
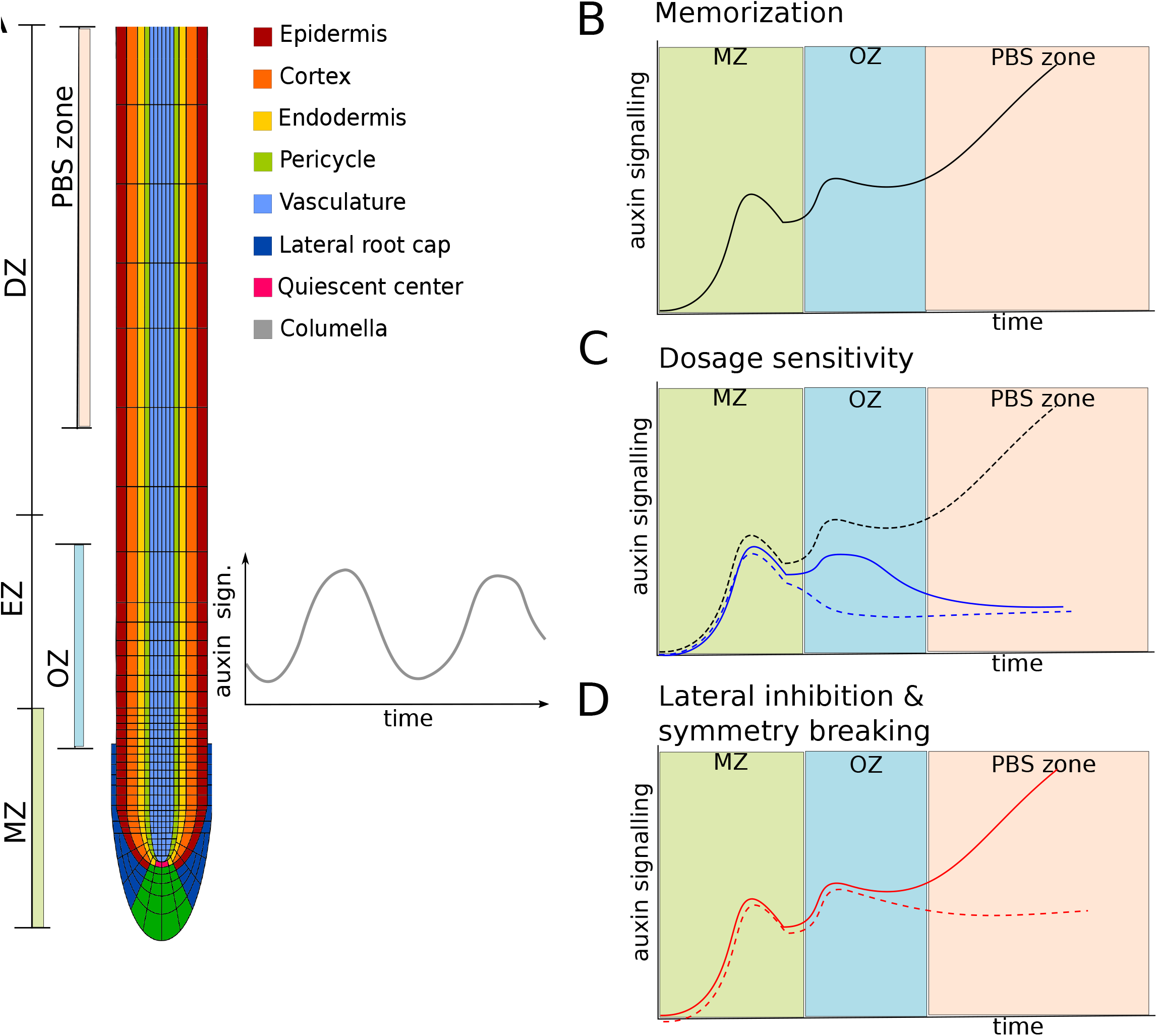
Auxin signalling characteristics during priming and prebranch site formation. A) Anatomy of the Arabidopsis root tip indicating the different cell types, with priming starting in the xylem pole vasculature and being transmitted to the pericycle, the developmental zones, with the meristem (MZ) where celll divisions occur, the elongation zone (EZ) where cells undergo vacuolar expansion and the differentiation zone (DZ) where cells acquire their terminal cell fate and overlapping with this the oscillation zone (OZ) where priming occurs and the prebranch site formation zone (PBS) where successful primings lead to stable PBS formation. B) Cells undergoing successful priming experience high auxin signalling levels in the tip of the meristem (MZ), auxin signalling subsequently becomes high again in the oscillation zone (OZ) during priming, and after a second decline becomes elevated for the third time leading to stable prebranch site formation (PBS) and thereby memorization of the transient priming signal. C) Cells in between priming events (blue dashed line), as well as cells undergoing priming under conditions with limited auxin availability (continuous blue line) experience lower auxin signalling levels in the OZ, and fail to establish a PBS. For comparison a line representing cells undergoing successful priming is added (dashed black line) D) Under normal conditions lateral inhibition between nearby PBS forming sites -both at the same and opposite vascular poles-leads to establishment and maintenance of one and repression the PBS forming site, preventing formation of multiple nearby PBS.

While these later stages of lateral root formation are relatively well studied, and the involved regulatory modules, generally centred around auxin signalling, and downstream processes have been well documented (Du and Scheres 2018; Lavenus et al. 2013; Santos Teixeira and Ten Tusscher 2019), we still have very limited knowledge on the early stages of lateral root formation. The very first step in lateral root formation has long remained enigmatic, with hypotheses for the mechanism underlying priming oscillations ranging from gravitropic bending (De Smet et al. 2007), and genetic oscillators (Moreno-Risueno et al. 2010), to the periodic shedding of the lateral root cap (Xuan et al. 2016). In a recent study we demonstrated that priming arises through a so-called reflux-and-growth mechanism involving the interplay of root tip auxin reflux and root growth dynamics (van den Berg et al. 2021). We revealed a critical role for tip driven root growth, which results in periodic variations in the sizes of cells arriving in the elongation domain, resulting in periodic variation in membrane surface area and hence passive auxin uptake capacity. The root tip auxin transport reflux loop results in an auxin loading domain specifically in this region. Together this gives rise to periodic oscillations in auxin loading and hence auxin levels in elongating cells.

However, it remains unclear how these priming events are subsequently transduced into the next developmental stages of lateral root formation. While priming occurs at the end of the meristem and start of the elongation zone, lateral root initiation occurs in the differentiation zone. This implies that the transient elevation in auxin that occurs during priming is somehow stably memorized and at a later stage sets in motion lateral root initiation. The observation that in case of successful priming auxin signalling levels first decline before rising again (Fig. 1B) (Xuan et al. 2015) supports the notion that this memorisation involves processes distinct from the initial priming. We have previously suggested that this requirement for two distinct processes, priming itself and its memorisation, may help resolve the debate on whether priming involves auxin signalling and gene expression versus auxin levels per se (Laskowski et al. 2008) but this remains to be tested.

Additionally, while priming occurs at both xylem poles in Arabidopsis and initially involves a larger area, wildtype lateral root formation typically occurs on a single side at a single location (Fig. 1D). In contrast, many mutants exist in which nearby lateral root formation, either at a single or opposing sides occur (Chang, Ramireddy, and Schmülling 2015; Fernandez et al. 2020; Hofhuis et al. 2013; Maule, Gaudioso-Pedraza, and Benitez-Alfonso 2013; Toyokura et al. 2019)(. These findings imply active lateral inhibition and symmetry breaking type mechanisms constraining lateral root formation. At least some of this inhibition appears to occur prior or during stable prebranch site formation (el-Showk et al. 2015; Toyokura et al. 2019). Finally, stable prebranch site formation is highly dosage sensitive, with low amplitude oscillations failing to result in prebranch site formation yet absence of auxin negative feedback mechanisms resulting in supernumerous prebranch sites that subsequently fail to develop into lateral roots (Perianez-Rodriguez et al. 2021; Xuan et al. 2015). The mechanism responsible for transducing bilateral priming events into unique, single sided stable prebranch sites in a dosage dependent manner has so-far remained elusive.

A major factor hampering progress in our understanding of these early stages of lateral root formation are their limited observability. Markers exist for later stages of lateral root development, with for example stably established founder cells being demarcated by GATA23 (GATA TRANSCRIPTION FACTOR 23) and MAKR4 (MEMBRANE-ASSOCIATED KINASE REGULATOR 4) expression, and ACR4 (ARABIDOPSIS CRINKLY 4) and ralfl34 (RALF LIKE 34) expression being involved in the formative divisions typical of lateral root initiation (De Rybel et al. 2010; De Smet et al. 2008; Murphy et al. 2016). After initiation, developing lateral roots can be distinguished based on typical root patterning transcription factors such as WOX5 (WUSCHEL-RELATED HOMEOBOX 5), SHR (SHORTROOT), SCR (SCARECROW) and the PLTs (PLETHORAs) (Du and Scheres 2017) but can also be microscopically discerned based on morphological characteristics (Malamy and Benfey 1997). In contrast, neither reporters nor anatomical markers exist enabling us to distinguish pericycle cells undergoing the changes from priming to initiation from those that do not. While single cell transcriptomics may eventually offer a means to distinguish subpopulations of xylem pole pericycle cells, given the low numbers of primed pericycle cells and the complex spatiotemporal patterning processes involved this has thusfar not been possible. As a consequence, it remains hard to experimentally determine how successfully primed cells and / or their surrounding cells differ from non-primed cells, and how these differences may contribute to the memorisation, and eventual symmetry breaking and lateral root initiation.

In this study we extend the model previously developed to decipher lateral root priming to investigate how priming may subsequently lead to stable prebranch site formation, a first prerequisite for lateral root initiation. We reverse engineer a hypothetical prebranch site formation mechanism that critically relies on temporal integration of the experienced priming signal with auxin induced upregulation of auxin signalling, and auxilliary roles for auxin induced upregulation of auxin transport and production. To determine the plausibility of the proposed PBS formation mechanism, we subsequently assess its compatibility with key characteristics of priming and PBS formation, lateral inhibition, symmetry breaking, and the spatial narrowing of the auxin signalling domain. Our hypothesized PBS formation mechanism suggests that from a very early stage onwards primed cells should become epigenetically distinct from their surrounding pericycle cells. Additionally, by revealing an essential role for priming driven upregulation of auxin signalling capacity in stable PBS formation, the proposed mechanism enables us to unite previously diverging viewpoints on the importance of auxin levels versus auxin signalling in lateral root prepatterning.

## Results

### Priming signal minimally maintained in pericycle cells

Recently we uncovered how root tip polar auxin transport produces an auxin loading zone at the start of the elongation zone (EZ), while root growth generates periodic variations in the size with which cells enter this zone, causing periodic variations in auxin loading potential particularly in narrow vasculature cells with high surface to volume ratios (van den Berg et al. 2021). Combined this generates the oscillations in vascular auxin levels that underly lateral root priming. To investigate how these auxin dynamics may lead to stable PBS formation we here use a root tip model with an extended differentiation zone (DZ) and highly regular division dynamics (see Methods). This enables us to follow auxin signalling dynamics over a prolonged spatiotemporal interval (Fig 2A, kymograph), and ensures regular, reproducible priming dynamics (Fig 2B) (movie 1). This regularity allows us for later figures to only display dynamics in the first, fifth, ninth and tenth cell of the first priming event shown in Figs 2B (arrows). To validate the robustness of our results, we confirmed that for staggered rather than parallel cell wall positioning similar priming dynamics occur (Suppl Fig 2, for details see Methods).

**Fig 2.**
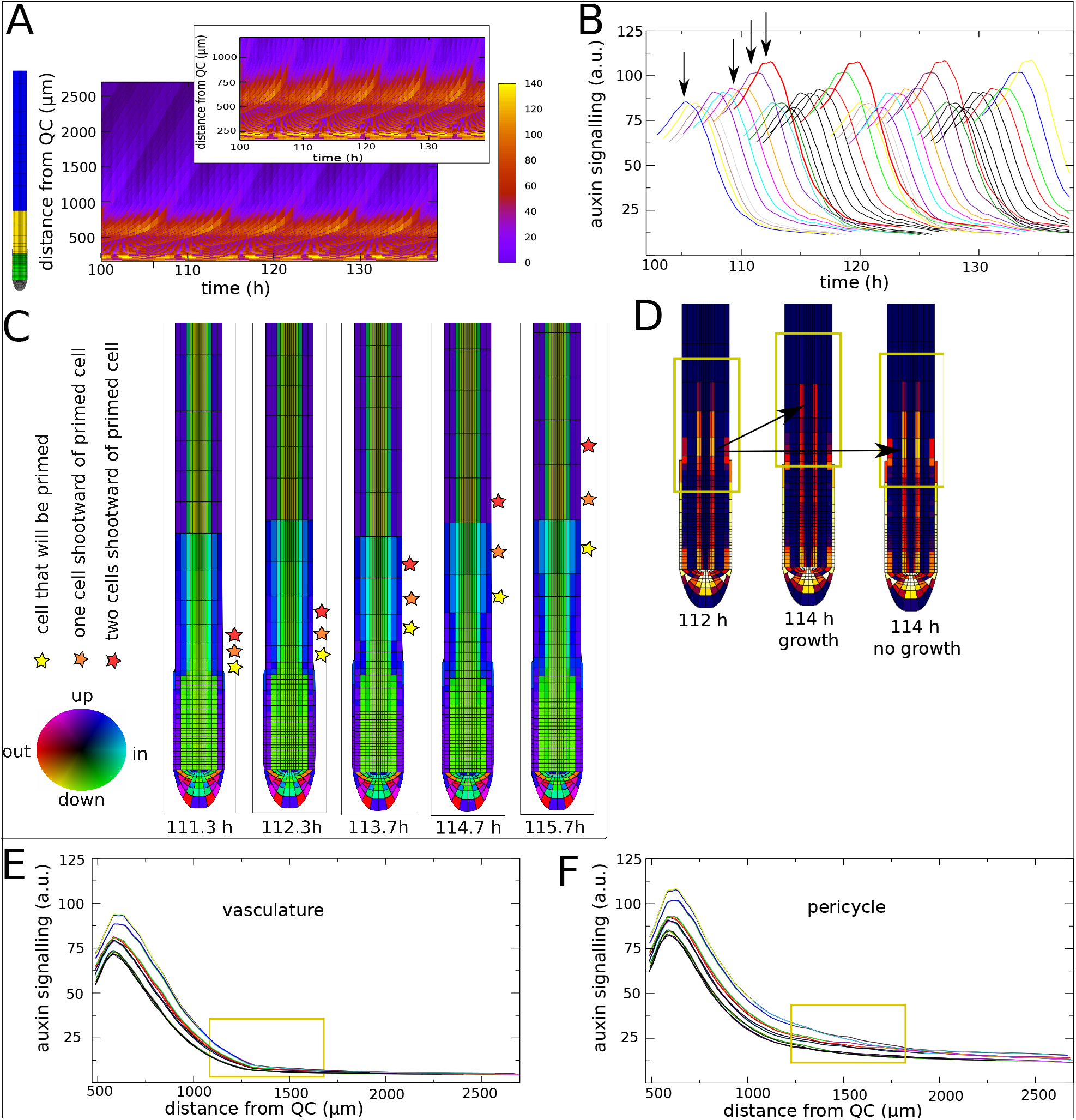
Generation and dissipation of the priming signal. A) Kymograph showing the spatiotemporal dynamics of auxin signalling in the pericycle over the timecourse of 5 priming events for the default settings of our root tip model. Snapshot on the left shows developmental zonation in the model root to enable interpretation of the distance from the root tip. Inset shows a kymograph for the same 5 priming events zoomed in on the lowermost parts of the root tip to enable comparison with our previous results (VandenBerg_2021). B) Auxin signalling dynamics as a function of time for a series of pericycle cells traced over time once they reach a distance of 500*μm* from the root tip. The traced cells correspond to the sequence of cells ranging from the first cell after the first priming event till the final primed cell of the fifth priming event shown in A, thus spanning a sequence of 4 priming events that each consist of 10 cells. C) Temporal sequence of auxin flux direction snapshots from the early (111.3 h) to late stages (115.7 h) of a single priming event. D) Auxin signalling dynamics in the lowermost parts of the root during early phases of priming (112 h), and at later stages of priming (114 h) in presense or absence of continued growth E) Auxin signalling dynamics as a function of distance for 40 vascular cells neighboring the 40 pericycle cells shown in B. F) Auxin signalling dynamics as a function of distance for the same pericycle cells as shown in B. In B, E and F line color is based on the order in which cells reach a distance of 500 *μm* from the root tip.

In Figure 2C we show a sequence of snapshots of auxin flux directions. We see that as EZ cells elongate epidermal and cortical fluxes reorient from predominantly upward to upward and inward, while endodermal fluxes change from downward to inward, together generating an auxin flux towards pericycle and vasculature cells. For cells arriving largest, this flux direction transition occurs closest to the start of the EZ (compare cells indicated with yellow star to those indicated with orange and red stars), consistent with the enhanced auxin loading potential we earlier uncovered for these cells. Still, while cell sizes keep increasing and once switched flux directions remain constant until cells enter the DZ, auxin levels in primed cells quickly dissipate (Fig 2A,B; movie 1). We hypothesized that this dissipation is due to the general decline in auxin levels that occurs as cells, due to growth and division of younger cells below them, become displaced to positions further away from the root tip. To investigate this we artificially stopped root growth after a priming event, resulting in auxin levels being maintained rather than dissipating (Fig 2D).

Fig. 2E and 2F show that while auxin oscillations and dissipation occur in both the vasculature and pericycle, in the pericycle the cells that receive the highest auxin levels are able to maintain their elevated auxin levels to a limited extent, while such maintenance is absent in the vasculature (yellow boxes). This can be understood from the changes in PIN patterns occuring when cells enter the DZ, with particularly endodermal and cortical PIN patterns becoming more apolar (for details see Methods). Indeed, from the auxin flux directions we observe that in the DZ the vasculature flux orients towards the pericycle, resulting in a passing on of remaining auxin to the pericycle. Additionally, endodermis and pericycle fluxes become oriented towards one another, giving rise to a recycling of auxin previously shown to be essential for maintaining and enhancing pericycle auxin levels (Laskowski_2008). Still, end EZ and DZ pericycle auxin signalling levels are far below those observed *in planta* during PBS formation.

### Direct positive feedback fails to differentially maintain auxin signalling in primed cells

In planta, auxin signalling levels in primed cells are observed to first decline and then rise again, suggesting a second, separate mechanism for stable prebranch site formation (Laskowski and Ten Tusscher 2017). Positive feedback would be a logical mechanism to amplify and maintain differences in auxin levels and/or signalling between cells. Indeed, auxin dependent induction of auxin importers such as AUX1 (AUXIN RESISTANT 1) and LAX3 (LIKE AUX1 3) (Laskowski et al. 2008), as well as a LEC2 (LEAVY COTELYDON 2) and FUS3 (FUSCA3) induced upregulation of YUCCA4 mediating auxin biosynthesis (Tang et al. 2017) have been implicated in early stage lateral root formation. The initial decline and subsequent rise of auxin signalling imply that these positive feedbacks need to become activated with a certain delay. Elevated cytokinin signalling in the early elongation zone could be a potential candidate for the initial suppression of these positive feedback mechanisms. For simplicity in our model we assume that additional LAX3 expression and YUCCA4 expression (we ignore the intermediary factors LEC2 and FUSCA3) can only occur once cells have reached a particular developmental stage (Fig 3A), which occurs at approximately 880 *μm* from the root tip, when the original priming signal is halfway its decline.

**Fig 3.**
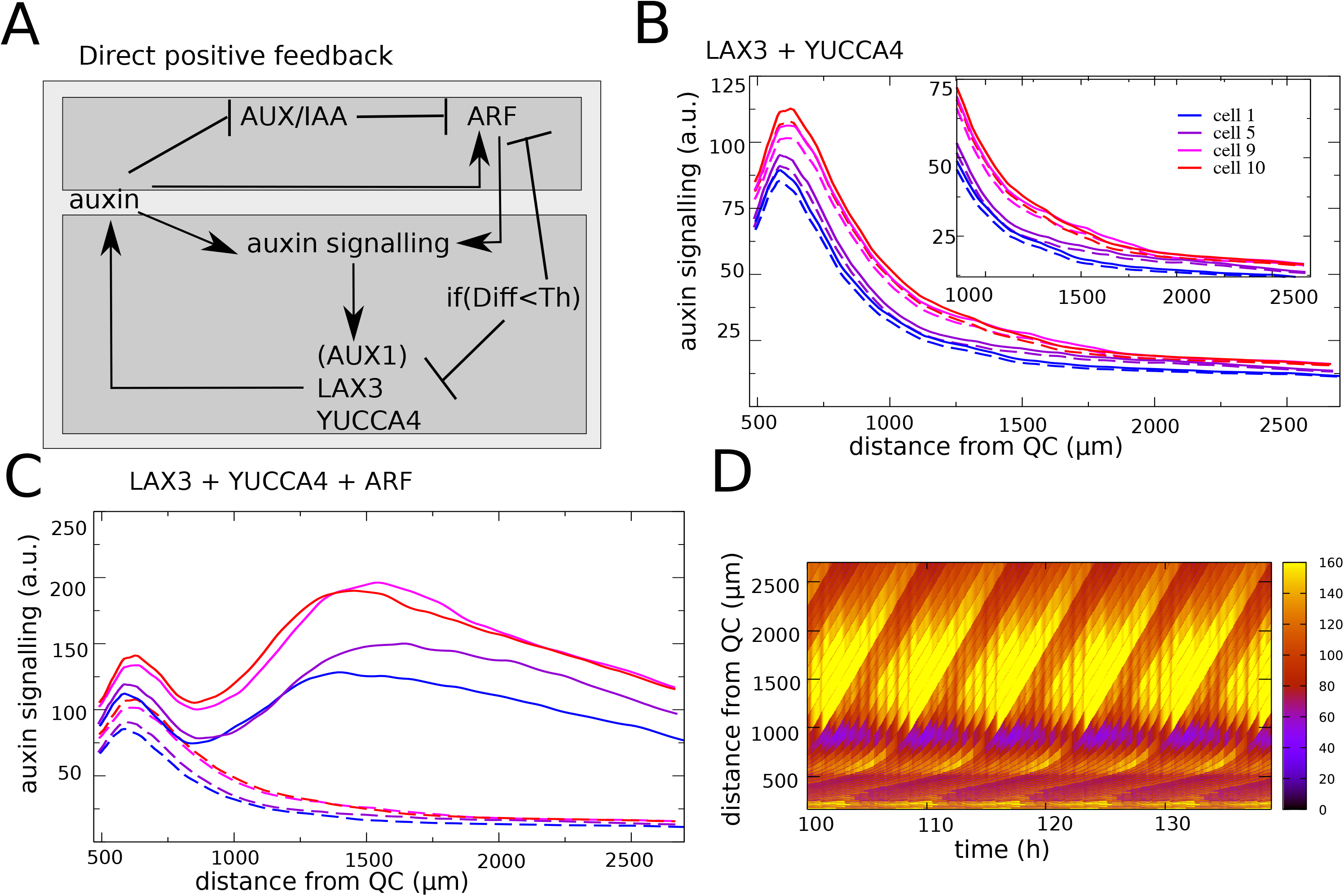
Positive feedback generates a secondary auxin signalling response. A) Overview of regulatory interactions added to generate positive feedback and enhance auxin signalling levels. A first feedback loop occurs between auxin signalling dependent expression of LAX3 and YUCCA4, a second feedback loop occurs between auxin signalling and ARF expression. Auxin signalling driven upregulation of ARF, LAX3 and YUCCA4 expression is repressed below a certain cellular differentiation level. AUX1 and LAX3 expression may occur in all cells, while YUCCA3 and ARF expression are restricted to occur in the elongation and differentiation zones to ensure activation in response to priming induced auxin elevation but not due to the high auxin levels in the meristem. Additionally, YUCCA3 expression is limited to the vasculature and pericycle based on experimental data (Tang_2017). B) Pericycle auxin signalling when only the LAX3 + YUCCA4 positive feedback loop is added. Auxin dynamics is shown for the first, fifth, ninth and tenth cell in a priming event. (maximum expression level of LAX3 and YUCCA4=100, Km =20). C) Pericycle auxin signalling when both feedback loops are added (maximum expression level LAX3 and YUCCA4=100, maximum expression ARF=300, Km=30). For comparison auxin signalling dynamics in absence of these feedbacks are shown (dashed lines). D) Kymograph pericycle auxin signalling corresponding to the settings in C.

Even when applying a low auxin induction threshold values and/or high maximum expression levels for LAX3 and YUCCA4, only a very limited secondary enhancement of auxin signalling levels was observed (Fig 3B, Suppl Fig 3A, B). Previously, it has been suggested that auxin signalling rather than auxin levels per se increase substantially as part of priming (Moreno-Risueno et al. 2010). We therefore hypothesized that enhancement of auxin signalling capacity could help in generating an increase in auxin signalling despite an overall decrease in auxin levels. We therefore speculated that ARF (AUXIN RESPONSE FACTOR) 5, 7, 19 known to play crucial roles during early stage lateral root formation (Du and Scheres 2018; Lavenus et al. 2013) may require transcriptional upregulation. Addition of an auxin-inducable generic ARF in our model, again gated to become expressed only beyond a specific cellular developmental stage (Fig 3A), indeed induces a significant secondary increase in auxin signalling levels (Fig. 3C). However, since auxin signalling ultimately depends on auxin availability, also in case of enhanced signalling capacity auxin signalling eventually declines. Additionally, although quantitatively different, primed and non-primed cells show qualitatively similar behavior in terms of this secondary increase in auxin signalling, something inconsistent with experimental results. This qualitative behavior is independent of the precise maximum level of ARF expression and its auxin induction threshold (Suppl Fig 3C,D). Finally, while additional LAX3 and induction of YUCCA4 expression quantitatively contribute to a secondary rise in auxin signalling, ARF induction alone is sufficient (Suppl Fig 3E).

### Temporal integration of priming signal robustly discerns primed cells

A recent study on phyllotaxis, the auxin-driven periodic patterning of leaves in the plant shoot apex, demonstrates that it is a temporally integrated rather than instantaneous auxin signal that drives the auxin-dependent transcription of developmental genes (Galvan-Ampudia et al. 2020). Furthermore the authors suggest that one likely mechanism for such temporal integration of auxin signalling is through epigenetic state changes. Inspired by this we decided to model the chromatin state of an at this stage not yet further specified gene, calling it EpiO, where low values indicate a closed and high values an open chromatin state. Experimental data indicate that auxin-dependent TIR1/AFB signalling, through the degradation of Aux/IAA and resulting liberation of ARF results in the recruitment of chromatin modifiers enhanching an open chromatin state (K. Lee, Park, and Seo 2017; Wu et al. 2015), while opposing factors contribute to a closed chromatin state (Fukaki, Taniguchi, and Tasaka 2006). Based on this we assume a basal closed chromatin state, with chromatin opening increasing with auxin signalling level. Additionally, closing of chromatin closing is assumed to decrease with both auxin signalling level as well as chromatin open state (see Methods). The latter reflects that an open chromatin state results in gene expression, which subsequently counteracts chromatin closing.

First we study EpiO dynamics in absence of additional LAX3, YUCCA4 or ARF expression. Comparing normalized auxin signalling with normalized EpiO dynamics we see that for sufficiently slow dynamics EpiO state is capable of temporally integrating auxin signalling, accumulating the smaller, instantaneous auxin signalling differences occurring at each time instance into a difference in chromatin open state that is significantly larger (see Suppl. Fig 4A,B for non-normalized auxin and EpiO dynamics). For our default parameter settings, normalized auxin signalling at the peak differs 25 a.u. (100 versus 75), whereas EpiO levels at the peak differ 54 a.u. (100 versus 46), a more than 2 fold increase in differences (Fig. 4B). Additionally, we also observe that EpiO differences are significantly longer maintained than auxin signalling differences. At a distance of 1000 microm from the root tip, auxin signalling differences have declined to 17 a.u. (45 versus 28), whereas EpiO levels still differ 44 a.u. (70 versus 26), a 2.5 fold increase in differences (Fig. 4B). This longer maintenance is promoted by high EpiO levels slowing down EpiO decay, representing transcriptional activity slowing down chromatin closing. Thus, particularly at the later stages that are relevant for the secondary increase of auxin signalling, EpiO levels are considerably more suited to differentiate primed from non-primed cells, and hence for selectively amplifying and memorizing auxin signalling in these cells.

**Figure 4.**
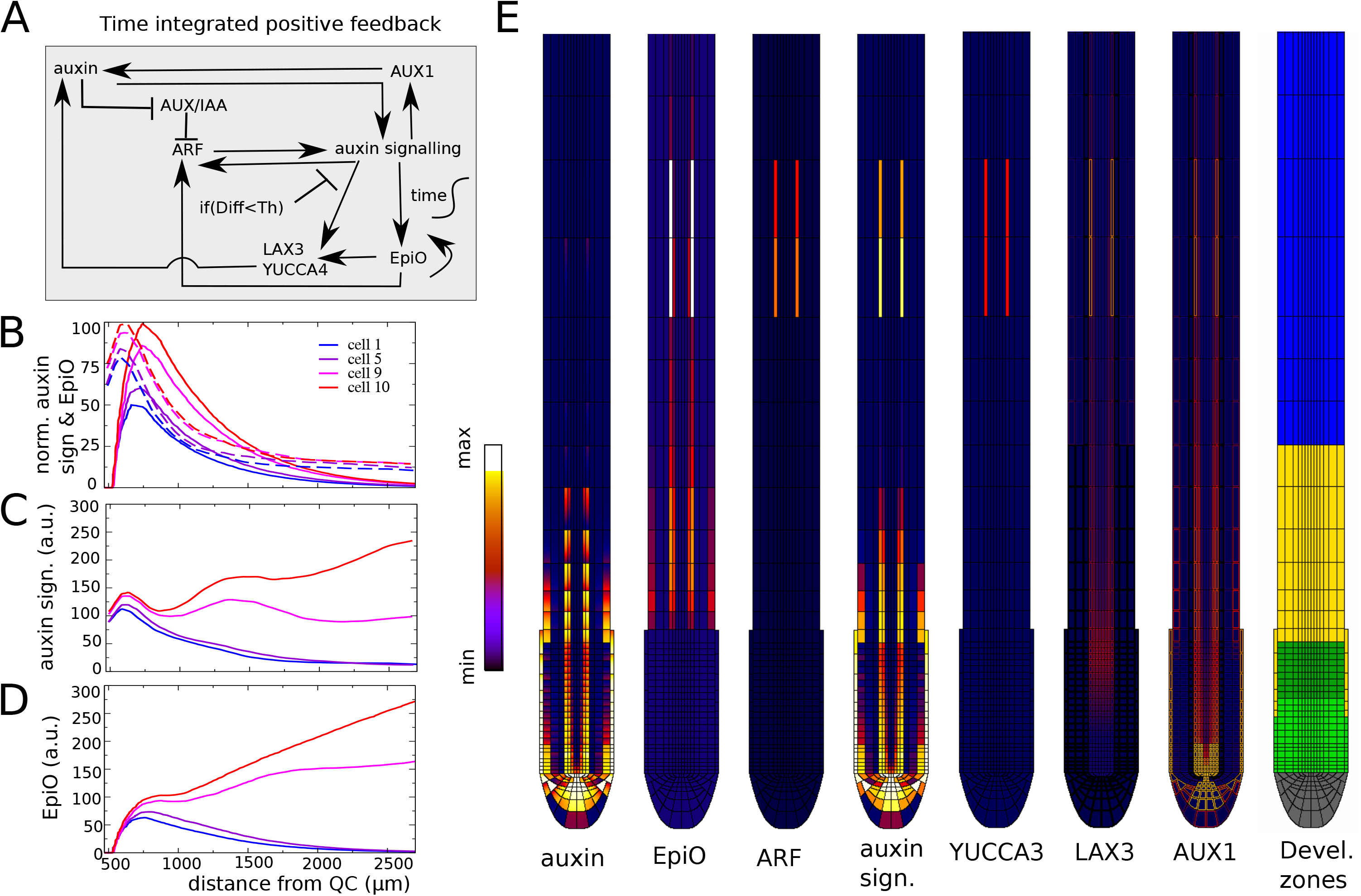
Time integrated positive feedback enables stable prebranch site formation. A) Overview of regulatory interactions in the model settings where positive feedback on ARF, LAX3 and YUCCA4 is gated by EpiO state. Again auxin signalling driven upregulation of ARF, LAX3 and YUCCA4 expression is repressed below a certain cellular differentiation level. B) Comparison of (normalized) auxin signalling (dashed lines) and EpiO (normal lines) dynamics for default parameter settings and in absence of positive feedback. C,D) Pericycle auxin signalling (C) and EpiO (D) dynamics as a function of distance in the combined EpiO+positive feedack model. E) Snapshot of auxin concentration, EpiO state, ARF, auxin signalling, YUCCA4 and LAX3 levels and developmental zonation patterns illustrating the induction of a high EpiO state resulting in high ARF, YUCCA4 and LAX3 expression and high auxin signalling levels in 2 pairs of cells that have undergone priming.

Precise buildup and maintenance of chromatin open state depends on parameter values. More rapid EpiO dynamics result in more rapid EpiO buildup and hence stronger amplification of auxin signalling differences, yet this occurs at the cost of maintenance (Supp Fig. 4C). In contrast, lowering the auxin signalling threshold for chromatin opening (EpiO buildup) enhances both maximum amplitude and maintenance of EpiO (Suppl Fig. 4D). In absence of data parameter values were chosen such that buildup and prolonged maintenance of EpiO signal is well supported.

### Temporal integration based positive feedback maintains auxin signalling in primed cells

Having established the suitability of an auxin-dependent chromatin open state in discerning primed from non-primed cells we now reinstate the previous positive feedbacks. We assume that epiO state gates the auxin-dependent expression of ARF, LAX3 and YUCCA4, allowing for their transcription only beyond a certain level of chromatin opening (see Methods). In case of LAX3, we assume epiO only gates part of its expression, e.g. through gating the expression of upstream LBD regulators that induces extra LAX3 expression (Lee et al. 2009), while other regulation of LAX3 is assumed independent of epiO. This latter aspect is based on the fact that LAX3 expression is not limited to primed cells and their neighborhood but occurs in a large region of the vasculature (el-Showk et al. 2015). Again, to insure a secondary, delayed increase in auxin signalling after priming, ARF, LAX3 and additional YUCCA3 expression only occur once cells have reached a particular developmental stage (Fig. 4A).

Figure 4C-E show that by using EpiO to gate LAX3, YUCCA4 and ARF expression, a much more selective enhancement of auxin signalling only in cells receiving high auxin levels during priming occurs. Additionally, because of the additional positive feedback now present between EpiO status and ARF expression, a stably maintained, continuous rising of auxin signalling level occurs consistent with experimental observations (Fig 4C). Finally, interestingly, for the applied parameter settings, auxin signalling in the cell experiencing the second highest auxin levels during priming also undergoes this continuous rise, albeit at lower levels. Stable prebranch site formation can be restricted to a single cell (albeit still on both sides of the root) by decreasing the maximum level (Suppl Fig 5A), or increasing the activation threshold (Suppl Fig 5C) of ARF expression while increasing the auxin signalling activation threshold for LAX3 and YUCCA4 had a negligible effect on auxin signalling dynamics (Suppl Fig 5E-F). Finally, while additional LAX3 and induction of YUCCA4 significantly contributes, ARF induction alone is again sufficient to drive a secondary rise in signalling (compare Fig 5C with Suppl Fig 5G).

**Fig 5.**
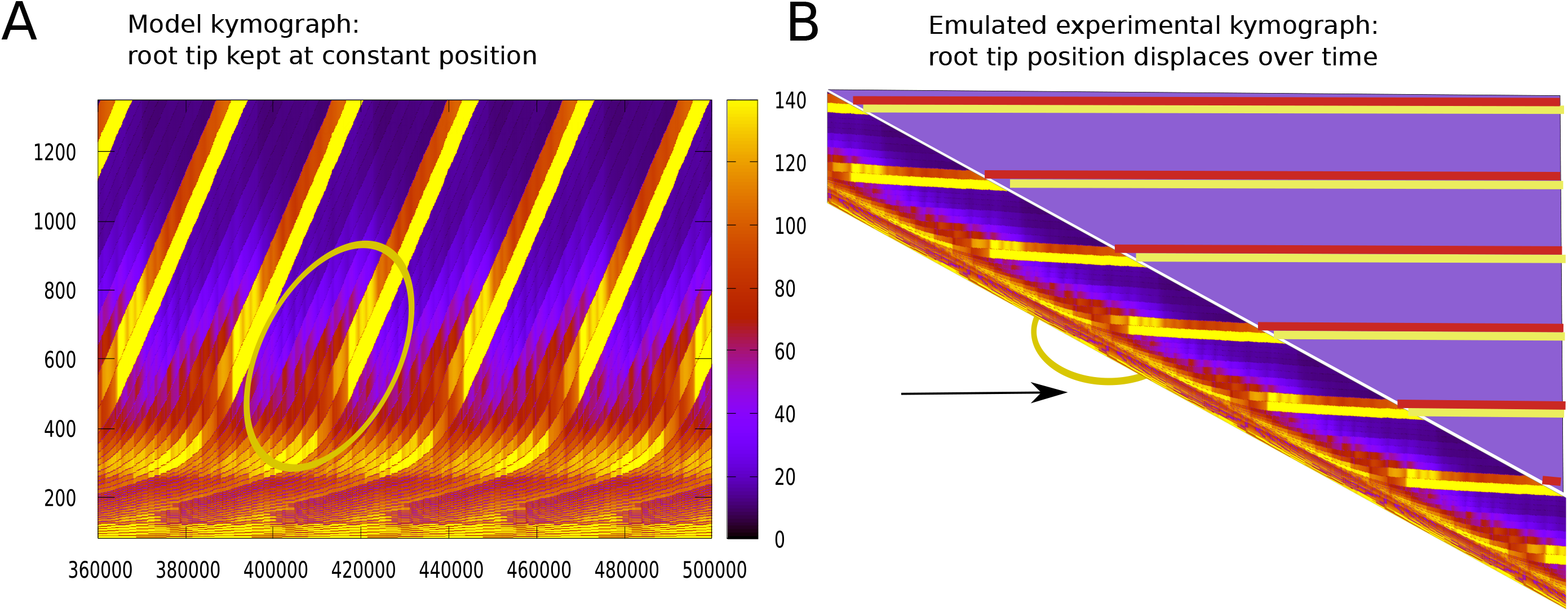
Narrowing and increase of auxin signalling during PBS formation. A Standard model kymograph for the model simulation shown in Figure 4 C-E. B) Kymograph emulating experimental kymographs. The kymograph was obtained by skewing the kymograph in A such that individual cells in the differentiation zone are maintained at a constant horizontal reflecting the absence of growth in this region. This automatically results in the continuous downward displacement of the lower parts of the root tip in which growth occurs. To further enhance the resemblance we artificially added what the kymograph would look like if in our model no culling of cells at the top of the simulation domain would occur but instead an increasingly large domain would be simulated as the root grows.

An important hallmark of priming and stable prebranch site formation is that while initially auxin signalling occurs in a relatively broad domain during and directly after priming, the auxin signalling domain subsequently narrows while signalling intensity increases, settling into a narrow auxin signalling peak demarcating a stable PBS (Moreno-Risueno et al. 2010). To investigate whether the PBS formation mechanism we propose here is consistent with this observed dynamics we look in detail at the spatiotemporal auxin signalling dynamics (Fig 5). To more clearly illustrate how the kymographs in our model are related to the auxin signalling space-time plots commonly applied in experimental papers on LR priming (Duan et al. 2021; Kircher and Schopfer 2018; Xuan et al. 2015, 2016), we show both our standard model kymograph in which root tip position is fixed at the base of the diagram (Fig 5A), and a transformation thereof in which root position moves downward through growth (Fig 5B). In our model, we simulate a constant sized root domain, culling cells as they approach the simulation domain boundary (see Methods), while experimentally as the root tip grows the size of the domain monitored increases in parallel. To further clarify the relation between our model results and experimental observations we added in shaded coloring what the kymograph would have looked like if the simulation domain were to grow. In both diagrams (inside yellow ovals) we observe that initially a group of ~5 cells experience an intermediate level of auxin elevation (red colors) and that over time this narrows down to 2 cells experiencing considerably higher auxin levels (yellow coloring), with over time a further narrowing as one of these cells decreases its auxin levles (orange coloring). Thus the PBS formation mechanism proposed here is consistent with experimentally observed spatial narrowing of auxin signalling.

### Lateral inhibition and symmetry breaking determine final PBS spacing

It has been previously shown that a significant fraction of priming events are not transduced into stable prebranch site formation but instead lead only to a transient increase in auxin signalling (Kircher and Schopfer 2018). Furthermore, in case of spatio-temporally nearby positioned priming events, a range of lateral inhibition type mechanisms has been uncovered to repress nearby lateral root formation. Described ACR4, PLETHORA, and plasmodesmatal effects on repressing nearby lateral root formation are only activated at or after the lateral root initiation stage (De Smet et al. 2008; Hofhuis et al. 2013; J Maule, Gaudioso-Pedraza, and Benitez-Alfonso 2013), while symmetry breaking due to auxin-dependent expression of AUX1 (el-Showk et al. 2015), the LBD16-TOLS2-RLK7-PUCHI pathway (Fig 6A) (Toyokura et al. 2019) and possibly also cytokinin signalling mediated repression (Chang, Ramireddy, and Schmülling 2015) are active from earlier stages onward. To investigate the compatibility of the PBS formation mechanism proposed here with early stage lateral inhibition we keep parameter settings as in Fig 4, such that a secondary PBS with lower auxin signalling levels is formed and next investigate how this secondary PBS can become repressed.

**Fig. 6.**
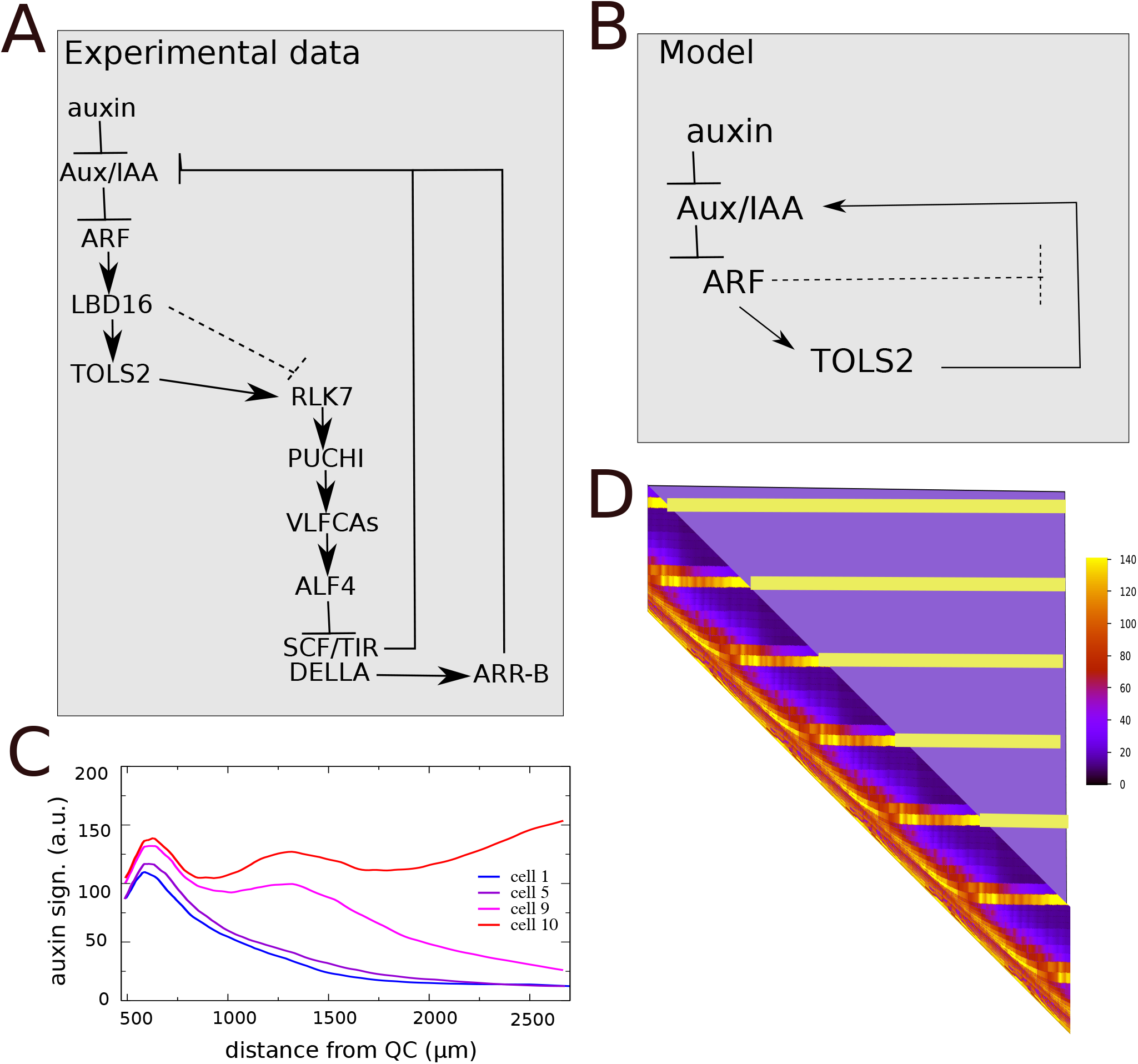
TOLS2 signalling mediated lateral inhibition. A) Layout of the network of experimentally uncovered regulatory interactions between LBD16 (LATERAL BOUNDARY DOMAINS 16), TOLS2 (TARGET OF LBD SIXTEEN 2), RLK7 (RECEPTOR LIKE PROTEIN KINASE 7), PUCHI, VLFCAs (very long chain fatty acids), ALF4 (ABERRANT LATERAL ROOT FORMATION PROTEIN 4), SCF/TIR, DELLA and Aux/IAA. For lateral inhibition to work, TOLS2 signalling should repress auxin signalling in neighboring cells, yet not in the cell itself, suggesting that a thusfar undiscovered interaction exist repressing RLK7 signalling in high auxin signalling cells (dashed interaction). B) Simplified TOLS2 network incorporated in the root tip model. C) Pericycle auxin signalling dynamics in the model complemented with TOLS2 signalling. D) Experimental style type kymograph of pericycle auxin signalling dynamics of the same model simulation.

Auxin-dependent AUX1 (and LAX3) expression is already part of our baseline model. Therefore next we added a simplified version of the LBD16-TOLS2-RLK7-PUCHI pathway to our model (Fig 6A,B). Based on experimental data showing that PUCHI results in enhanced production of VLFCAs (Trinh et al. 2019), which have been shown to upregulate of ALF4 (Shang et al. 2016), a repressor of SCF/TIR and DELLA (Bagchi et al. 2018), this ultimately results in the upregulation of Aux/IAA in cells neighboring the high auxin and high LBD16 expressing cells. Fig 6 shows that for sufficiently strong TOLS2 signalling inclusion of this mechanism enables the rootward stronger PBS to repress the shootward weaker PB (Suppl Fig 6), consistent with experimental observations (Toyokura et al. 2019). Note that because of the mutual repression between forming PBS, also in the remaining stable PBS overall auxin signalling levels have become somewhat lower.

In planta, symmetry breaking between left and right PBS is expected to be initiated through stochastic differences in the sizes of left and right xylem pole pericycle cells, their expression of auxin importers and exporters as well as root bending. Here, we incorporated a limited level (10%) of initial asymmetry in AUX1 expression in our model to explore whether symmetry breaking would occur. We find that while a significant enhancement of the initial asymmetry occurs, with the shootward, weaker PBS being repressed more rapidly and the remaining, rootward PBS obtaining lower auxin signalling levels at the low AUX1 side (Suppl Fig 7A) no full repression of PBS at the weaker AUX1 side occurred for our default parameter settings. If we additionally changed the Km of AUX1 from 75 to 85, making it less auxin sensitive, we do observe a complete repression of PBS at the weaker AUX1 side (Fig 7A), where an initial small asymmetry during double sided priming is translated into single sided PBS formation (Fig 7B). Full symmetry breaking could also be obtained by elevating the Km of LAX3 or enhancing the effect of TOLS2 signalling (Suppl Fig 7B,C), supporting the idea that these inhibitory mechanisms act in an additive manner. Additionally, similar results were obtained for imposing an initial asymmetry in e.g LAX3 instead of AUX1 (Suppl Fig 7D). Finally, our model standard incorporates a weak, radial distance dependent, concentration difference dependent auxin flow between corresponding positions in the left and right side of the root (see Methods) to take into account the 3D nature of actual plant roots. To investigate the importance of this 3D auxin flow coupling we either removed or enhance this coupling (Suppl Fig8A,B), demonstrating that this reduces or enhances respectively the symmetry breaking between opposite sided PBS, supporting the relevance of 3D auxin flows.

**Fig. 7.**
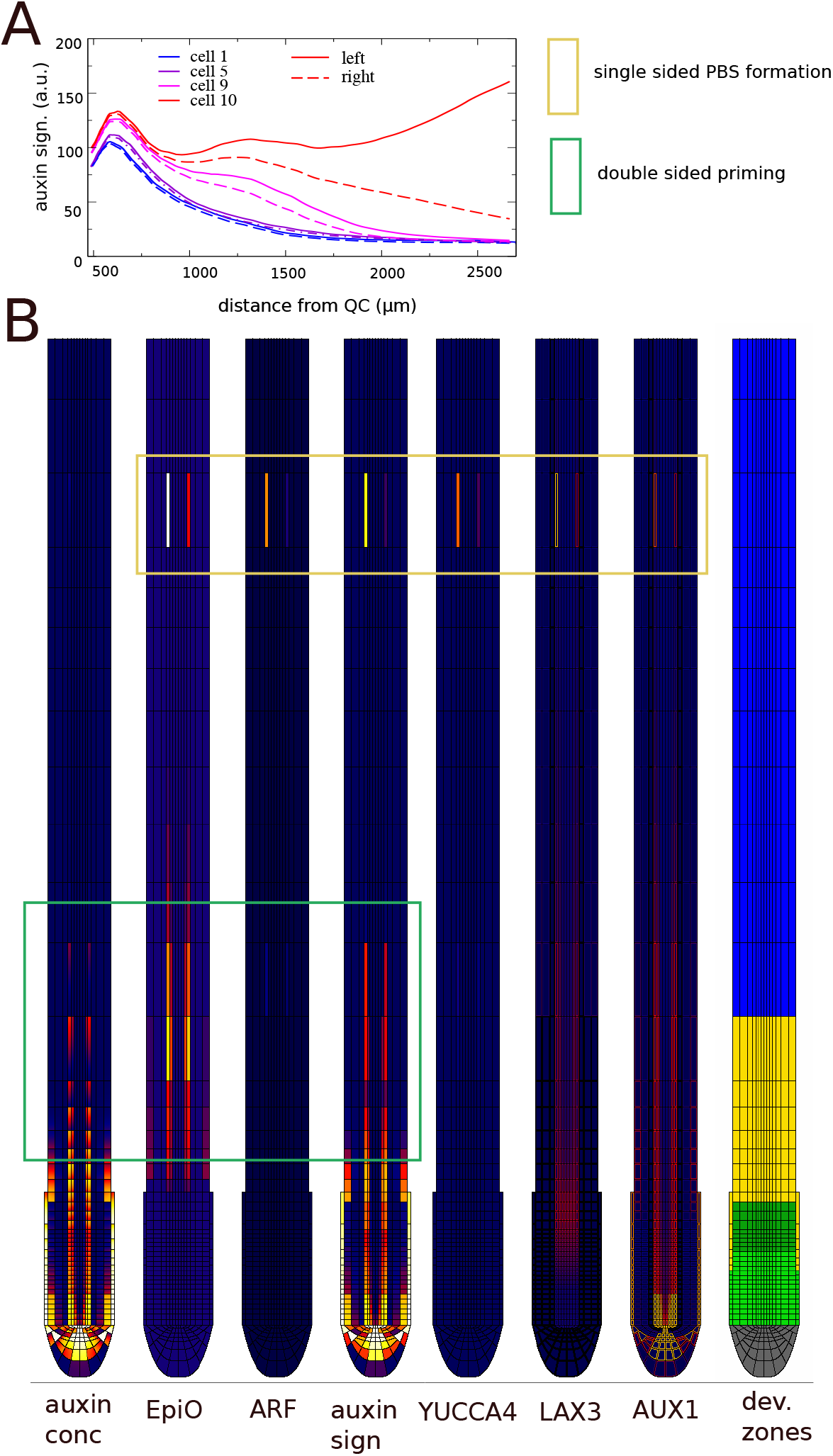
PBS symmetry breaking. A) Auxin signalling dynamics in left (black) and right (red) xylem pole pericycle cells. B) Snapshot of auxin, EpiO, free ARF, auxin signalling, YUCCA4 expression, LAX3 and AUX1 membrane levels and developmental zonation. Bottom part of the snapshots shows symmetric priming (green box) and top part shows asymmetric prebranch site formation (yellow box).

### Dosage dependence and importance of auxin homeostasis for priming

In planta, due to stochasticity in root bending, cell growth, division, gene expression etcetera, there is inherent variability in auxin levels between different priming events. In case overall auxin levels are high, most priming events result in sufficiently high auxin signalling levels to lead to stable prebranch site and lateral root formation. In contrast, if auxin availability is lowered through e.g. mutants reducing the production of the auxin precursor IBA in the lateral root cap, average priming amplitude is decreased, and only the higher amplitude subset of priming events results in stable prebranch site formation (Xuan et al. 2015). In our deterministic model this variability between priming events is absent and we simply test whether a lowering of auxin levels results in failure of stable prebranch site formation. Figure 8A shows that indeed a lowering of root auxin content reduces the auxin levels during the initial priming event and subsequently prevents stable prebranch site formation.

**Fig. 8.**
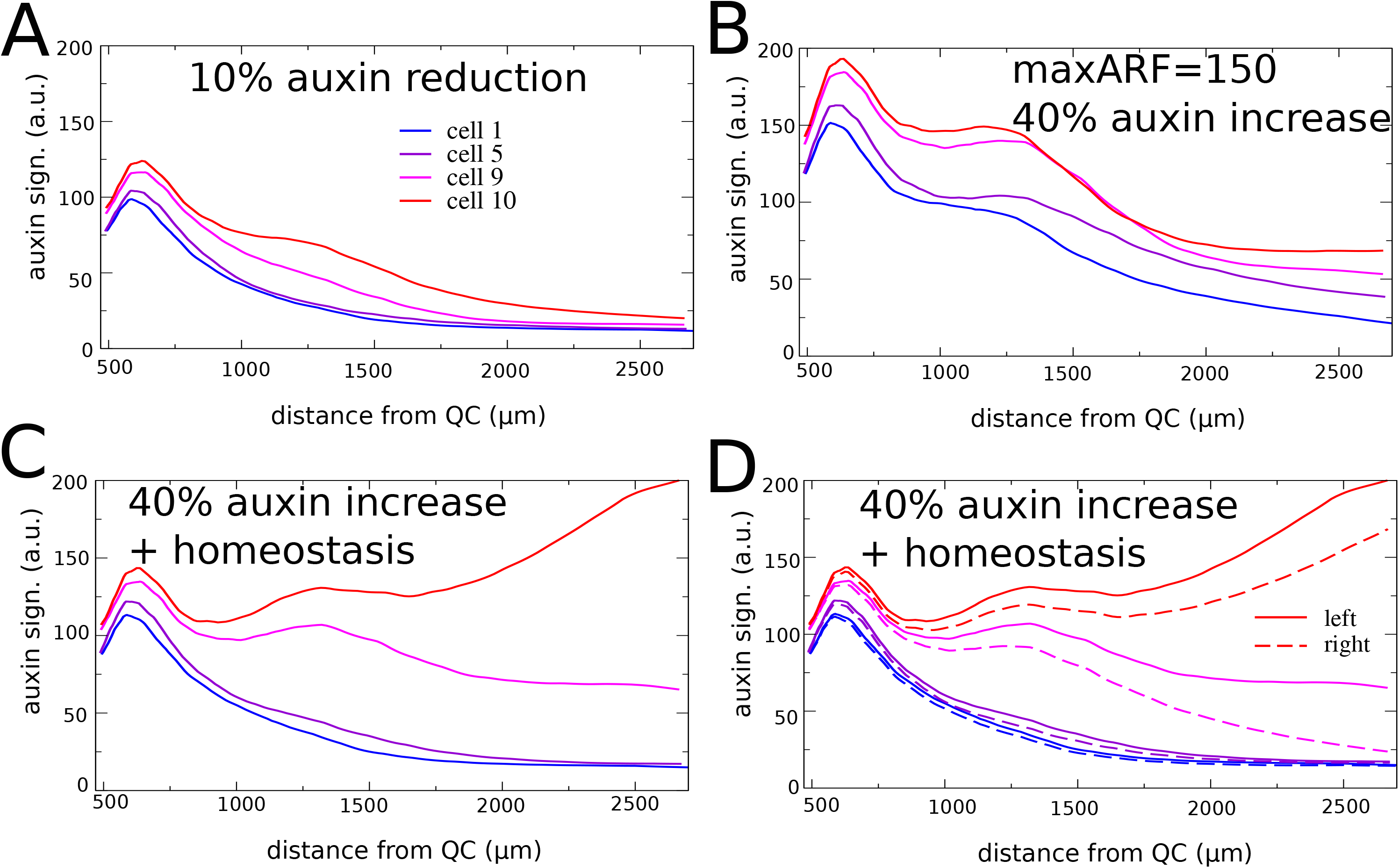
Auxin dosage effects priming effectiveness. A) A reduction of 10% in root tip auxin content reduces priming amplitude and abolishes stable prebranch site formation. B) A 50% reduction of maximum ARF levels combined with a 40% increase in root tip auxin generates transient secondary elevations in auxin signalling in most cells. C,D) Combining a 40% increase in auxin inflow with an auxin homeostasis mechanisms restores the selective secondary auxin signalling elevation in 2 consecutive cells and the subsequent lateral inhibition between them (C), yet does not fully restore left-right symmetry breaking (D)

In addition to low auxin availability being detrimental for lateral root formation, it was recently shown that also elevated auxin levels may significantly reduce lateral root numbers. Mutations in either ARF7 or IAA18/POTENT (auxin insensitive mutant) where shown to result in elevated auxin levels, an increase in prebranch site formation and subsequent failure of lateral root formation (Perianez-Rodriguez et al. 2021). These mutations where interpreted as defects in the root clock, an alternative mechanism proposed for priming. In contrast, we argue that in absence of an experimentally reported mechanism for cell-autonomous gene expression oscillations and considering the clear effects on auxin levels, these mutants should be considered as defects in root auxin homeostasis. To investigate this hypothesis, we simulate a single potent or arf7 mutant in our model through a decreased expression of the generic ARF (50%), assuming that while ARF7 is non functional, other ARFs like ARF19 are still functioning, as well as an increased auxin availability as a result of this mutation (influx increased by 40%). Figure 8B shows that most cells rather than only those experiencing the highest initial auxin signalling levels generate a secondary response, consistent with the experimental observation of prolific PBS formation in potent/arf7 mutants. Interestingly, this secondary response is more modest and in later stages dissipates, with even the highest initial auxin signalling experiencing cells not able to mount the continuous increase in auxin signalling observed for wildtype conditions. Again, this is consistent with observations for the potent/arf7 mutants which after prolific PBS formation fail in subsequent lateral root development (Perianez-Rodriguez et al. 2021). Our results indicate that less auxin signalling capacity can not be simply compensated for by more auxin. Since auxin itself spreads out due to transport, while auxin signalling is cell bound, instead more cells undergo a less effective secondary auxin signalling response . To further support this possible role of excess auxin underlying the potent/arf7 phenotype next added both the extra auxin as well as a simplified, ARF-dependent auxin degradation to our simulation with normal, wildtype ARF levels (see Methods). Under these conditions the secondary increase in auxin signalling indeed remains restricted to the two highest auxin experiencing cells at each root side, with subsequent lateral inhibition between these two sites (Fig 8C). Still, not only is lateral inhibition a bit slowed at the high auxin side as compared to earlier (Fig 8C), also left right symmetry breaking is not fully restored (Fig 8D). Our results thus support that auxin homeostasis is critical to prevent precocious yet non-effective PBS formation yet also indicate that in planta auxin homeostasis is spatiotemporally more refined than currently incorporated in our model.

## Discussion

In the current work we extended on our previous efforts to elucidate the earliest steps of lateral root formation through the use of multi-scale modeling. Previously we elucidated a so-called reflux-and-growth mechanism for lateral root priming (van den Berg et al. 2021). We demonstrated how the root tip auxin reflux loop establishes an auxin loading domain at the start of the elongation zone which combined with growth driven periodic variations in cell sizes and hence auxin loading capacity result in semi-periodic oscillations in cellular auxin levels. Still, in isolation reflux-and-growth priming results in only transient increases in auxin levels that dissipate as cells are displaced further from the root tip. Thus, this still leaves open the question of how succesfull priming is transduced into stable prebranch site formation as characterized by a stable peak of auxin signalling (Moreno-Risueno et al. 2010).

Here, using a computational reverse engineering approach, we established essential ingredients for a secondary rise in auxin signalling leading to stable prebranch site formation. First, we established that simply incorporating positive feedbacks through adding auxin-dependent expression of auxin importing and biosynthesizing proteins is insufficient both in specifically targeting primed cells and generating a significant secondary rise in auxin signalling. We demonstrated that an auxin-dependent upregulation of ARF genes enables an increase in auxin signalling capacity that counteracts the dissipation of auxin levels in primed cells. Our results thus indicated a primary role for upregulation of auxin signalling (ARF) and more auxilliary roles for auxin production and transport (YUCCA4, LAX3) in generating a secondary auxin signalling response. Additionally, we found that responding to a temporally integrated auxin level, which we here assumed to arise from auxin dependent chromatin opening, rather than instantaneous auxin signalling, enables for a robust distinction between primed and non-primed cells. Combined, these two processes enable specifically those cells experiencing the highest auxin levels to mount a secondary responses that increases over time, in agreement with experimental observations (Toyokura et al. 2019; Xuan et al. 2015). The mechanism proposed here bears close resemblance to observations made on phyllotaxis, in which a temporally integrated auxin signal was shown to induce the transcriptional response of cells (Galvan-Ampudia et al. 2020), and to recent findings on the formation of fractal floral architectures where a key role was identified for stable memorization of transient LEAFY expression (Azpeitia et al. 2021).

The mechanism for PBS formation identified here reproduces the experimentally observed spatial narrowing and simultaneous increase in auxin signalling as a stable PBS is formed. Additionally, it is consistent with previously identified mechanisms for lateral inhibition and symmetry breaking that translate a double-celled double-sided priming into a single-celled single-sided PBS. In the current model, only auxin-dependent AUX1 and LAX3 and the ARF-LBD-TOLS2-RLK7-PUCHI mechanisms for competition between neighboring PBS were integrated, whereas in planta, an even larger repertoire of inhibitory processes takes place. Recently the TOLS2-RLK7-PUCHI and the GLV6/GLV10-RGI peptide signalling pathway were identified as highly convergent lateral inhibitory processes (Jourquin et al. 2022) and one could interpret the mechanism implemented here rather as a proxy for the combination of these two mechanisms. While the mechanisms incorporated in our model are active at early stages, during the translation from priming signal to stable prebranch sites, other mechanisms are thought to play a role at later stages. For example, ACR4 limits periclinal divisions to a limited number of cells during lateral root initiation (De Smet et al. 2008), while expression of the PLETHORA 3,5 and 7 transcription factors and normal plasmodesmatal dynamics in developing lateral root primordia inhibit formation of nearby primordia (Hofhuis et al. 2013; J Maule, Gaudioso-Pedraza, and Benitez-Alfonso 2013). This is consistent with our model findings that depending on the strength of the AUX1/LAX3/PUCHI mechanisms relative to the auxin signalling levels, early stage lateral inhibition may be complete or not, explaining the need for later stage inhibitory mechanisms.

We believe that the mechanism proposed here also offers a reconciliation of the dispute on whether priming and prebranch site formation involve an elevation in auxin concentration levels, or rather involves an increase in auxin signalling (Moreno-Risueno et al. 2010). Our model results suggest that while the initial priming involves an elevation in auxin concentration, for it to become effectively translated into a stable prebranch site a subsequent elevation in auxin signaling is essential. This corresponds to the previously suggested distinction between priming signal and memory formation (Laskowski and Ten Tusscher 2017). Furthermore, assuming that ARF7 is the, or one of the, ARFs upregulated after priming, our model also explains the earlier observed oscillations in ARF7 expression (Moreno-Risueno et al. 2010). Moreno-Risueno et al. previously reported that out-of-phase addition of auxin does not result in ARF7 expression or additional prebranch site formation, arguing that thus auxin alone is insufficient for prebranch site formation. Our current results suggest that the auxin application was too limited or short to change epigenetic state and enable ARF7 expression and PBS formation. Additionally, the lateral inhibitory mechanisms active during the translation from priming to prebranch site formation may well explain the failure of additional prebranch site formation. Finally it was recently shown that in both ARF7 and IAA18(potent) mutants, auxin levels and the number of prebranch sites increased, while subsequent lateral root development was blocked (Perianez-Rodriguez et al. 2021). Interpreting these mutants as auxin homeostasis rather than clock mutants, and simulating this through addition of excess auxin, our model provided an explanation for these seemingly counterintuitive results. While the excess of auxin enabled ARF7 activation and PBS formation in most cells, explaining supernumerous PBS formation, individually these cells reached lower auxin signalling levels as compared to when only primed cells undergo PBS formation. This explains the halting of further lateral root development. Adding a naive, auxin signalling level dependent degradation of auxin as homeostasis could not fully restore normal PBS formation, indicating that in planta homeostasis is more complex. Indeed, if auxin degradation were directly downstream of ARF7, preferential degradation at auxin signalling maxima would occur whereas in planta this may well be directed towards neighboring competing cells.

In conclusion, our results suggest that while the priming signal consists of an elevation in auxin concentration, its conversion to a PBS involves a time-integrated measuring of this auxin and a subsequent translation into increased auxin signalling. Assuming this time integration has an epigenetic basis, our results suggest that epigenetic marks on auxin signalling factors involved in PBS formation could serve as an early marker distinguishing these from other pericycle cells. Our work demonstrates that auxin levels and signalling capacity can not be traded against each other, instead more auxin with less signalling capacity results in many cells undergoing non-effective PBS formation. We thus clearly defined separate essential roles for auxin and auxin signalling. Sufficient auxin arising in a subset of cells during priming is needed to activate an enhanced auxin signalling capacity, with this triggered auxin signalling subsequently enabling further LR development.

## Methods

### Root anatomy, cell types and developmental zones

We simulate root growth, auxin and gene expression dynamics on a realistic root tip topology, similar to previous studies (van den Berg et al. 2016, 2021; Cruz-Ramírez et al. 2012; Di Mambro et al. 2017; Salvi et al. 2020). However, in contrast to these earlier studies, in order to follow what happens over time to primed sets of pericycle cells and track whether these develop into stable prebranch sites, we model a significantly larger part of the differentiation zone (Suppl Fig 1A). This enables us to follow these cells as over time due to growth and division of new cells their distance from the root tip increases and they occupy a position further and further into the differentiation zone.

Our model includes from inside to outside epidermis, cortex, endodermis, pericycle and vascular cell types. Additionally, in the root tip it incorporates lateral root cap, columella and quiescent center cell types (Suppl Fig 1A). In addition to cell types, developmental zones are included. From the root tip shootwards we include the meristem proper, in which cell divisions occur, the transition zone, in which cells no longer divide but still undergo slow cytoplasmic growth, the elongation zone, in which cells undergo rapid vacuolar expansion, and the differentiation zone, in which cells attain their final differentiation characteristics (Suppl Fig 1A). For simplicity, the location and size of these zones are superimposed in this model.

| parameter | meaning | default value | alternative values | units |
| --- | --- | --- | --- | --- |
| **In all models** |  |  |  |  |
| $p_{auxin,baseline}$ | Basal rate of auxin production | 0.0013 | Not varied | $[\text{s}^{-1}]$ |
| $celltype\ factor$ | Cell type dependent auxin production rate enhancement factor | 100-50-30-1<br>QC, top<br>columella, initials<br>– rest columella –<br>lateral root cap - rest | Not varied | dimensionless |
| *yucca4factor* | YUCCA4<br>expression based<br>auxin production<br>rate enhancement<br>factor | 0.1625 | Not varied | dimensionless |
| $d_{auxin}$ | Auxin degradation<br>rate | 0.00007975 | Not varied | $s^{-1}$ |
| $eff_{basal}$ | Basal auxin efflux<br>rate | 0.95 | Not varied | $s^{-1}$ |
| $p_{PIN}$ | PIN mediated<br>auxin efflux rate | 0.2 | Not varied | $[\ ]^{-1}s^{-1}$ |
| PIN | Constant cellular<br>PIN expression<br>level | 100 | Not varied | $[\ ]$ |
| $inf_{pas}$ | Passive auxin<br>influx rate | 2.5 | Not varied | $s^{-1}$ |
| $p_{AUX/LAX}$ | AUX/LAX<br>mediated auxin<br>influx rate | 0.07 | Not varied | $[\ ]^{-1}s^{-1}$ |
| $D_{cell}$ | Intracellular auxin<br>diffusion rate | 600 | Not varied | $\mu m^2 s^{-1}$ |
| $D_{wall}$ | Intra-apoplastic<br>auxin diffusion rate | 40 | Not varied | $\mu m^2 s^{-1}$ |
| $auxin_{ext}$ | Concentration of auxin in the first row of cells of the not-explicitly simulated rest of the plant | 0.96 | 0.912 Fig 8A<br>1.344 Fig 8B-D | [] |
| $p_{AUX1}$ | Maximum AUX1 production rate | 0.015 | Not varied | [s<sup>-1</sup>] |
| $K_{m,AUX1}$ | AUX1 half-saturation constant | 85 | 75 Fig 7A | [] |
| $d_{AUX1}$ | AUX1 degradation rate | 0.00015 | Not varied | s<sup>-1</sup> |
| $p_{Diff}$ | Maximum differentiation factor production rate | 0.01 | Not varied | [s<sup>-1</sup>] |
| $d_{Diff}$ | Differentiation factor degradation rate | 0.0001 | Not varied | s<sup>-1</sup> |
| $avgdivduration$ | Duration of division cell cycle | 9 | Not varied | h |
| $avgexpduration$ | Duration of cell expansion | 7 | Not varied | h |

| Added in the direct positive feedback model |  |  |  |  |
| --- | --- | --- | --- | --- |
| $p_{LAX3,basal}$ | Baseline LAX3 production rate | 0.004 | Not varied | [s<sup>-1</sup>] |
| $p_{LAX3,auxin}$ | Maximum auxin-dependent LAX3 production rate | 0.01 | 0.005 SupplFig. 3B | [s<sup>-1</sup>] |
| $K_{m,LAX3}$ | LAX3 half-saturation constant | 30 -direct model<br>75 -time integrated model | 50 SupplFig. 3A | [] |
| $d_{LAX3}$ | LAX3 degradation rate | 0.0001 | Not varied | s<sup>-1</sup> |
| $p_{YUCCA4}$ | Maximum YUCCA4 production rate | 0.01 | 0.005 SupplFig. 3B | [s<sup>-1</sup>] |
| $K_{m,YUCCA4}$ | YUCCA4 half-saturation constant | 30 -direct model<br>60 -time integrated model | 50 SupplFig. 3A<br>80 SupplFig. 5E<br>40 SupplFig. 5F | [] |
| $d_{YUCCA4}$ | YUCCA4 degradation rate | 0.0001 | Not varied | s<sup>-1</sup> |
| $unbind$ | AUX/IAA ARF dissociation rate | 0.001 | Not varied | s<sup>-1</sup> |
| $bind$ | AUX/IAA ARF association rate | 0.05 | Not varied | [<sup>-1</sup>s<sup>-1</sup>] |
| $pIAA$ | IAA production rate | 0.01 | Not varied | [s<sup>-1</sup>] |
| $d_{IAAbasal}$ | Basal IAA degradation rate | 0.0001 | Not varied | $s^{-1}$ |
| $d_{IAATIR1}$ | Auxin-TIR1/AFB mediated IAA degradation rate | 0.001 | Not varied | $s^{-1}$ |
| $p_{ARFbasal}$ | Basal ARF production rate | 0.001 | Not varied | $[s^{-1}]$ |
| $p_{ARFauxin}$ | Maximum auxin induced ARF production rate | 0.03 | 0.02 SupplFig. 3C<br>0.0225 SupplFig. 5A<br>0.035 SupplFig. 5B | $[s^{-1}]$ |
| $K_{m,ARF}$ | ARF half-saturation constant | 30 -direct model<br>30 -time integrated model | 75 SupplFig. 3D<br>50 SupplFig. 5C<br>20 SupplFig. 5D | $[]$ |
| $d_{ARF}$ | ARF degradation rate | 0.0001 | Not varied | $s^{-1}$ |
| **Added in the time integrated positive feedback model** |  |  |  |  |
| $open_{EpiO}$ | Maximum EpiO opening rate | 0.012 | 0.0168 SupplFig. 4C | $[s^{-1}]$ |
| $close_{EpiO}$ | Maximum EpiO closing rate | 0.00012 | 0.000168 SupplFig. 4C | $s^{-1}$ |
| $frac_{min}$ | Minimum constant fraction of EpiO<br>closing rate | 0.2 | Not varied | dimensionless |
| $K_{m,EpiO,1}$ | Half-saturation<br>constant for auxin<br>dependent<br>promotion of<br>opening en<br>inhibition of<br>closing of EpiO | 95 | 75 SupplFig. 4D | [] |
| $K_{m,EpiO,2}$ | Half-saturation<br>constant for EpiO<br>dependent<br>inhibition of<br>closing of EpiO | 75 | Not varied | [] |
| $p_{vasc}$ | Vascular<br>production rate of<br>LAX3 | 0.004 | Not varied | dimensionless |
| $p_{priming}$ | Priming related<br>production rate of<br>LAX3 | 0.012 | Not varied | dimensionless |
| $K_{m,LAX3,2}$ | LAX3 half-saturation constant<br>for priming related<br>part | 60 | 80 SupplFig. 5E<br>40 SupplFig. 5F<br>75 SupplFig. 7B | [] |
| $K_{m,trans}$ | Half-saturation constant for EpiO dependent gated transcription of LAX3, YUCCA4 and ARF | 85 | Not varied | [] |
| **Added for lateral inhibition** |  |  |  |  |
| $p_{TOLS2}$ | Maximum TOLS2 production rate | 0.1 | 0.03 SupplFig 6B | [s<sup>-1</sup>] |
| $K_{m,TOLS2}$ | TOLS2 half-saturation constant | 60 | Not varied | [] |
| $d_{TOLS2}$ | TOLS2 degradation rate | 0.0001 | Not varied | s<sup>-1</sup> |
| $D_{TOLS2}$ | TOLS2 diffusion rate | 0.04 | Not varied | $\mu m^2 s^{-1}$ |
| $p_{IAA,TOLS2}$ | TOLS2 mediated additional IAA production | 2.5 | 1.5 SupplFig 6A<br>3.5 SupplFig 7C | dimensionless |
| $K_{M,IAA,1}$ | Half-saturation constant for TOLS2 mediated IAA upregulation | 50 | Not varied | [] |
| $K_{M,IAA,2}$ | Half-saturation constant for ARF mediated<br>inhibition of IAA<br>upregulation | 60 | Not varied | [] |

### Auxin dynamics

Our root topology is laid out on a grid. Individual grid points belonging to the root tissue either correspond to the inside of a cell, a cell membrane or a wall. Except for the cells in the lowermost, curved part of the root tip, cells are modeled as rectangles of grid points. Cells have a cell type specific width, whereas cell height is a function of growth stage and developmental zone, similar to our previous models (van den Berg et al. 2021; Salvi et al. 2020). Because of this grid level subcellular resolution, our model enables the simulation of intracellular and intra-apoplastic auxin diffusion and the gradients this may result in.

Intracellular auxin dynamics are described using the following partial differential equation,

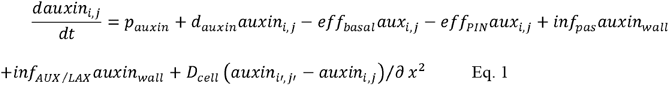

with *p_auxin_* auxin production, *d_auxin_* auxin degradation, *eff_basal_* basal ABCB mediated auxin efflux, *eff_PIN_* PIN-mediated auxin efflux, *inf_pas_* passive auxin influx, *inf_AUX_/_LAX_* AUX/LAX mediated influx and *D_cell_* intracellular auxin diffusion.

For auxin production *p_auxin_*, in addition to a baseline production rate, *p_auxin,baseline_*, we incorporate the experimentally observed higher production of auxin around the QC and in the columella, as well as the high production of auxin precursors in the lateral root cap (Suppl Fig 1B). For this we multiply the baseline production rate with a parameter *celltypefactor*, which equals 100 for the QC, top columella layer and vascular initials, 50 for the lower columella layers and 30 for the lateral root cap, for all other cell types celltypefactor is set to 1 (see Table 1). Additionally, based on its reported relevance for very early stages of lateral root development (Tang_2016), we incorporate a YUCCA4 expression dependent auxin production, which through the parameter *yucca*4*factor* is translated into an increase of auxin production rate. Parametrization is done such that at a maximum YUCCA4 expression level of 100, auxin production rate increases 16.25-fold (Table 1). Combined this gives rise to the following expression for auxin production:

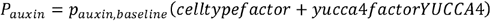

Since auxin production is applied per grid point, we finally normalize this auxin production relative to cell height to ensure that overall auxin production of a cell is not a function of cell height, by multiplying the above term with 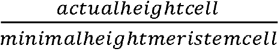.

For PIN-mediated auxin efflux *eff_PIN_*, we incorporate a cell and zone type PIN prepattern defining the maximum expression levels and polarity pattern (Suppl Fig 1C). Actual membrane PIN levels are defined as the product of these membrane grid-level prepattern levels and cellular PIN expression level, which in this model for simplicity we assumed constant (at a level of 100):

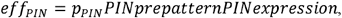

with *p_PIN_* the rate of PIN mediated transport.

Similarly, for AUX/LAX-mediated auxin influx *inf_AUX/LAX_*, we incorporate cell and zone type specific prepatterns for AUX1 and LAX3, (Suppl Fig 1D,E), while actual membrane AUX1 and LAX3 levels are defined as the product of prepattern level and gene expression level. AUX1 expression is standard auxin dependent, for model extensions auxin signalling dependent LAX3 expression is added (see below). Overall active influx is the sum of AUX1 and LAX3 mediated influx: 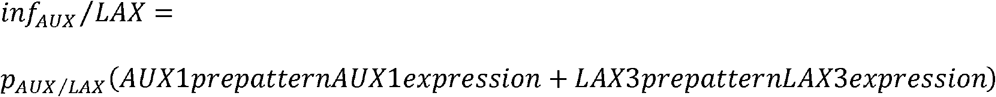, with *P_AUX/LAX_* the rate of AUX/LAX mediated transport.

Expression of AUX1 is auxin dependent in all model settings (see below), in the baseline model LAX3 expression is set constant at a level of 40, in the direct and time-integrated feedback model settings it is auxin dependent (see below).

Values for the parameters discussed above and in the next sections can be found in Table 1.

### Boundary conditions

Similar to previous root tip models developed by us as well as others (van den Berg et al. 2021; Grieneisen et al. 2007; Mähönen et al. 2014), we simulate auxin exchange with the not-explicitly modeled rest of the plant. For this we model an inflow of auxin into the walls above the topmost vascular, pericycle and endodermal cells, and an outflow of auxin from the walls above the topmost epidermal and cortical cells. The inflow term consists of a constant, low, external auxin concentration (*auxin_ext_*) multiplied by the efflux (basal and PIN mediated) of not-explicitly simulated cells one layer above the topmost simulated cells, and which is taken equal to the efflux of these topmost simulated cells. The outflow term consists of the local wall auxin concentration multiplied by 10% of the influx (passive and AUX/LAX) of not explicitly simulated cells, which is taken equal to the influx of the topmost simulated cells.

### Gene expression dynamics

Gene expression dynamics are modeled on a cellular level, using ordinary differential equations.

For simplicity we do not distinguish separate transcription and translation dynamics, but only describe resulting protein dynamics.

#### Baseline model

In our baseline model, following our earlier work in (van den Berg et al. 2021), we incorporate the auxin dependent expression of the auxin importer AUX1:

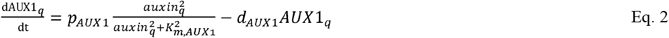

with *p*_AUX1_ production rate, *K*_m,AUX1_ the half-saturation constant of auxin dependent transcription, and *d*_AUX1_ degradation rate.

To describe the differentiation dynamics cells undergo upon entering first the elongation and subsequently the differentiation zone we describe the dynamics of a generalized transcription factor as follows:

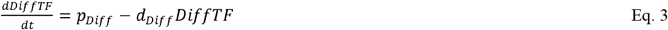

with *p_Diff_* the production rate of this differentation factor, which is set to zero as long as cells are in the meristematic zone and obtains a non-zero value upon elongation zone entry, and *d_Diff_* the degradation rate of this differentiation factor which in absence of ongoing differentiation (i.e. if cells were to revert to the meristematic zone due to changes in hormone or gene expression levels) allows for dedifferentiation.

#### Direct positive feedback model

As a next step, we extended the model with additional positive feedback regulations known or suggested to be involved in the earliest stages of lateral root formation.

First, also the auxin importers LAX3 is known to have auxin-dependent expression (Fig. 3A). We thus incorporated:

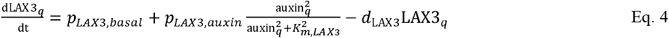

with *p*_*LAX*3,*basal*_ a basal production rate, *p*_*LAX*3,*auxin*_ the maximum auxin-dependent additional production rate, *K*_m,LAX3_ the half-saturation constant of auxin dependent transcription, and *d*_LAX3_ degradation rate.

Second, in early lateral root formation, pericycle specific expression of the transcription factors LEC2 and FUS3 has been shown to induce the auxin biosynthesizing enzyme YUCCA4 (Tang et al. 2017). Additionally, expression of FUS3, and indirectly also LEC2 is auxin-dependent (Gazzarrini et al. 2004; Horstman et al. 2017). Combined this gives rise to another positive feedback loop (Fig 3A) In our model we simplified this through incorporating an auxin-dependent expression of YUCCA4 specifically in the pericycle:

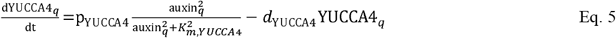

with *p*_YUCCA4_ production rate, *K*_m,YUCCA4_ the half-saturation constant of auxin dependent transcription, and *d*_YUCCA4_ degradation rate.

Third, it has been suggested that not only auxin levels, but also auxin signalling contributes to the early stages of lateral root formation (Moreno-Risueno et al. 2010). We thus assume that in addition to the default expressed ARFs (not modeled explicitly), auxin-elevation may cause additional ARF expression in elongation and differentiation zone pericycle cells, which may subsequently contribute to auxin-dependent gene expression (Fig 3A).

For ARF-TIR1-IAA-auxin dynamics we write:

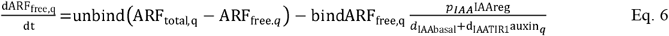

Whereas for ARF expression we write:

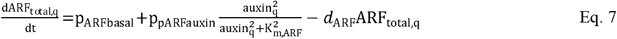

with *unbind* the dissociation rate of the ARF-AUX/IAA complex, *bind* the association rate between ARF and AUX/IAA, *pIAA* the AUX/IAA production rate, *d*_*IAAbasal*_ the baseline AUX/IAA degradation rate, *dIAATIR*1 the auxin-dependent TIR1/AFB induced extra degradation of AUX/IAA*p_ARFbasal_* the baseline ARF production rate, *p_ARFauxin_* the auxin-dependent ARF production rate *K_m,ARF_* the auxin-level at which auxin-dependent ARF production is half-maximal, and *d_ARF_* the ARF degradation rate.*IAAreg*, which is here set constant at 1 enables for the incorporation of additional regulatory effects impacting IAA levels, which we will use in a later section when incorporating lateral inhibition signalling.

To ensure that a secondary rise in auxin signalling occurs, auxin dependent increases in LAX3, YUCCA4 and ARF expression are assumed only to occur beyond a cellular differentiation level of 80 (i.e. the differentiation factor described in Eq 3 should be >80) (Fig 3A).

In the model settings incorporating the induction of ARF expression, to take into account the additional auxin signalling this results in we replace in the equations for AUX1, LAX3, YUC3 and ARF itself subsequently auxin_*q*_ by *auxsign*_*q*_ which is defined as:

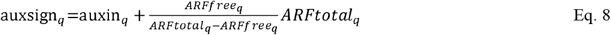

The rationale behind this equation is as follows. In the baseline model we did not explicitly model AUX/IAA ARF dependent auxin signalling but instead made gene expression a direct function of auxin levels. In the positive feedback models we now need to add an enhanced auxin signalling capacity through the additional expression of ARF. Normally one would translate auxin levels into the corresponding free ARF levels, however to be able to also keep working with the baseline auxin signalling formulated simply in terms of auxin concentration we here take an opposite approach and translate free ARF levels into a corresponding auxin level through taking the ratio between free and bound ARF and subsequently multiply this by total ARF expression as a measure of auxin signalling potential.

#### Time integrated positive feedback model

As a next model extension, we incorporate a variable *EpiO* representing an auxin dependent chromatin open state which through its slow dynamics enables time-integrated tracking of the auxin signalling experienced by cells:

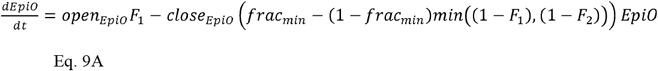

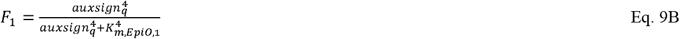

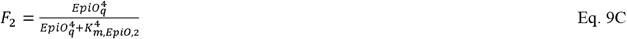

where *open_EpiO_* is the maximum rate of chromatin opening*close_EpiO_* is the maximum rate of chromatin closing, and *frac_min_* is the minimum fraction of the chromatin closing rate that is always effective. Chromatin opening rate is a saturating function *F*_1_ of auxin signalling, reaching its half maximum rate at an auxin signalling level of *K*_*M,EpiO*,1_. Chromatin closing is inhibited both by auxin signalling ((1 – *F*_1_)) and chromatin open state ((1 – *F*_2_)), assuming that beyond a certain threshold level chromatin open state leads to transcriptional activity and transcription limits chromatin closing.

This chromatin open state gates in a non-linear manner the auxin signalling dependent transcription of LAX3, YUCCA4 and ARF:

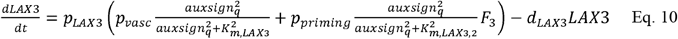

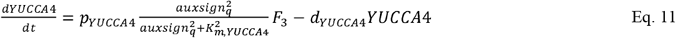

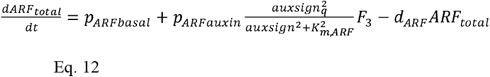

with

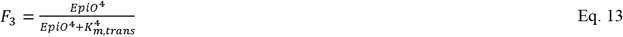

with *K_m,trans_* the EpiO level at which half-maximum auxin-dependent transcription is possible.

The non-linear function *F*_3_ implements the assumption that for effective transcription a sufficiently open chromatin structure is required. As before, the auxin signalling dependent induction of LAX3, YUCCA4 and ARF is represssed below a differentiation level of 80.

Note that while for YUCCA4 and ARF the gene expression equations are the same as before except for the EpiO dependent gating of their auxin induction, for LAX3 we used a slightly adjusted equation compared to before. Here, we split up the auxin-dependent gene expression into a part *P_vasc_* representing general vascular expression that is not related to lateral root priming and hence independent of EpiO gating and a part *p_priming_* that is related to priming and lateral root formation and hence is EpiO gated.

#### Lateral inhibition

To incorporate the ARF-LBD-TOLS2-RLK7-PUCHI lateral inhibition pathway (Fig 6A) in our model in a simplified manner (Fig. 6B) we make production of the TOLS2 signalling peptide directly dependent on free ARF levels and model its transport as simple cell to cell diffusion:

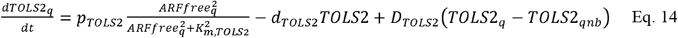

with *p*_*TOLS*2_ the TOLS2 production rate, *K*_*m,TOLS*2_ the TOLS2 half-saturation constant, *d*_*TOLS*2_ the TOLS2 degradation rate and *D*_*TOLS*2_ the TOLS2 diffusion rate.

To model the repressive effect of TOLS2 signalling on auxin signalling through elevating IAA levels, as well as the protective effect of the cells own ARF levels against this inhibition of auxin signalling to create lateral inhibition we replace the previously constant *IAAreg* = 1 in the free ARF equation with:

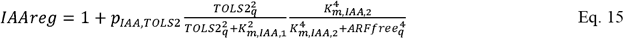

with *P*_*IAA,TOLS*2_ the additional TOLS2 mediated IAA production *K*_*m,IAA*,1_ the half-saturation constant for TOLS2 mediated IAA upregulation and *K*_*M,IAA*,2_ the half-saturation constant for free ARF mediated repression of IAA upregulation.

### Cellular dynamics

#### Zonation

As stated above, we superimpose locations and boundaries of the different developmental zones. Cells with for their lower cell boundary are at a distance of 514 *μm* or less from the root tip are part of the meristem, cells at a distance between 364 and 514*μm* from the root tip are part of the transition zone, the distalmost region of the meristem in which cells still grow cytoplasmatically but not longer undergo divisions.

For the faster maturing lateral root cap the meristematic zone ends at 314*μm* microm from the root tip. After this cells enter the elongation zone, and when their differentiation level exceeds a threshold transition to the terminal differentiation zone occurs.

#### Cell growth, division and expansion

In both the meristem and transition zone, cells undergo slow cytoplasmic growth. Identical to our previous models, cell growth, division and expansion are only implemented in the rectangular shaped cells positioned above the ending of the curved part of the lateral root cap and columella. Growth of the QC, columella and lower lateral root cap cells is thus ignored in our model. Since cells are modeled on a discrete grid, cells can only growth through the periodic, discrete addition of a row of grid points.

Previously we simply calculated the time till the next addition of a grid row to a cell as the inverse of the cellular growth rate, with the latter being the product of the per micrometer growth rate (see below) and the discrete, grid-based height of the cell. As a consequence, each time a grid row was added and the discrete height of the cell increased by one also a discrete jump in overall cellular growth rate occurred, whereas within this time interval growth rate stayed equal. While this effect is minor when cells in different cell files have the same height and positioning, this becomes problematic if say one big cell is neighbored by two cells half its height. Under these conditions, because the larger cell will overall grow more rapidly it will add grid rows and achieve a speedup in growth rate at earlier time points. This will cause it to become out of sync and start sliding relative to its neighbors, whereas on the per micrometer basis the one big and two small cells should grow equally fast. To minimize this unrealistic sliding and ensure a continuos rather than abruptly changing cellular growth rate we now endow cells also with a continuous valued cell length that is continuously updated based on a per micrometer growth rate. When this continuous valued cell length exceeds the integer, grid-based cell length by a value of 1 a new row of grid points is added to the cell and the integer cell length counter is updated.

Upon doubling their original cell size, cells in the meristematic zone divide, with daughter cells inheriting the PIN and AUX/LAX prepattern levels and polarity patterns of their mother cells. As cells at later stages enter the elongation and differentiation zone, cellular PIN and AUX/LAX prepatterns are adjusted to the zone specific patterns.

In the elongation zone, cells undergo rapid vacuolar expansion. Again this is implemented by the periodic addition of a row of grid points to increase cell length. As growth now arises from an increase in vacuolar volume rather than an increase in cytoplasmic volume nu dilution of protein levels is assumed to occur.

The per micrometer growth rates are computed from the duration of the division cell cycle or expansion duration (Table 1) respectively. For example, as cells need to double in size prior to division, solving 2*L* = *Le^rt^* for the given division duration *t* = *cellcycle* gives us the growth rate *r* for cell division related growth.

#### Cell differentiation

Upon entering the elongation zone, cells also start their initial differentiation. We model this through a generalized differentiation factor Diff undergoing a constant rate of differentiation *p_Diff_* as well as differentiation decay *d_Diff_*. The latter is incorporated to enable dedifferentiation of partly differentiated cells upon a perturbation that changes the identity of the developmental zone in which the cell is located (i.e. reversal from elongation zone to meristem due to e.g. elevation of PLETHORA levels).

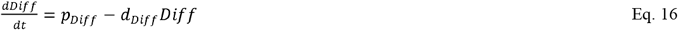

As cells exceed a threshold Th_Diff_ > 85 they cease elongation and enter the zone of terminal differentiation where further length increase ceases.

#### Semi-homogeneous cellular division rates

Recently we demonstrated that lateral root priming arises from the synergy between root tip cell division dynamics and auxin transport (van den Berg et al. 2021). In this previous study, to be able to closely compare model predictions to experimental data, we strived for maximally realistic root tip cell division dynamics, incorporating a gradient of division rates from QC via stem cells to transit amplifying cells as well as differences in cell size, division rates and zonation between vasculature and other cells. These detailed division dynamics resulted in realistic, semi-regular priming dynamics. Here, to study the translation of priming into prebranch site formation, we need to be able to discern transients in model dynamics from variations in priming dynamics between individual priming events in order to establish when stable prebranch site formation occurs. Therefore, we minimized variation in priming dynamics in our simulations through ignoring differences in cell division rates between QC, stem cells and TA cells or cell types within the meristem. To avoid artificial synchronized division of the entire meristem, deterministic position based small variations in division rates were applied. This was done as follows:

~~~
if(mean_j%3==0)
        divisionrate=average division rate
else if(mean_j%3==1)
        division rate =average division rate +5%
else if(mean_j%3==2)
        division rate= average division rate −5%
~~~

where mean_j is the longitudinal coordinate of the center of the cell after its origination from the division of the mother cell (i.e. it is determined once for each cell, not constantly updated).

Only in a subset of simulations (Fig 2E,F Suppl Fig 2) we incorporated cell height differences in the meristem (vasculature 12*μm*, epidermis and cortex 10*μm*, pericycle and endodermis 8*μm*), as well as differences in the onset of the transition zone between different cell types (as done previously in (van den Berg et al. 2021)) to obtain more realistic staggered cell patterns.

#### Constant sized finite simulation domain

In our model we make use of a constant, finite sized simulation domain. For the simulations used in this study, with a spatial integration step of *Δx* = 2*μm*, the simulation domain measures 141 times 1516 gridpoints or 282 times 3032*μm*.

Within this domain we keep the root tip at a fixed location in the bottom of the simulation domain. Hence, as a result of formation, division and growth of younger cells below them older cells are displaced towards the top of the simulation domain. If the wall above the apical membrane of a cell reaches the topmost position of the simulation domain this cell is removed from the simulation.

Additionally, to mimick the shedding of lateral root cap cells, the topmost lateral root cap cells are removed when the basal membrane of that cell exceeds the postion where other non-lateral root cap cells leave the meristem and enter the elongation zone (at 314*μm* from the root tip).

### Approximating 3D tissue architecture

In our simulations we model a two-dimensional longitudinal cross-section of the Arabidopsis root tip. Given that the Arabidopsis root tip, and particularly the vasculature, is not perfectly radially symmetric, the orientation of this cross-section was choosen such that it encompasses the for priming essential xylem poles and neighboring pericycle.

Still, by taking such a 2D approach, the radial auxin fluxes that occur in the 3D plant tissue are ignored, whereas in a previous modeling study taking radial instead of longitudinal cross-sections these have been previously shown to play an important role in the symmetry breaking between lateral root prebranch/initiation sites (el-Showk et al. 2015). Therefore, to at least partially emulate the 3D nature of true plant roots we incorporated radial auxin flows between laterally symmetric positions in the root.

In the default model settings we only incorporated a passive auxin exchange between laterally symmetric wall grid points. For this we added the following term to the auxin differential equations for the wall grid points:

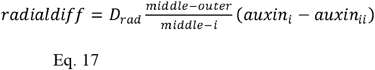

with *i* the coordinate of the left and *ii* the coordinate of the laterally symmetric right position, *D_rad_* the radial diffusion rate, *middle* the coordinate of the root tip middle and *outer* the coordinate of the leftmost wall of the root tip. Through incorporating 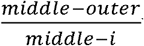 we inversely scale the rate of passive diffusion between laterally symmetric positions with their radial distance. All simuations except the one shown in Suppl Fig 8A incorporate these radial flows.

In the simulation in Suppl Fig 8B, in addition to the above described passive auxin flow between laterally symmetric wall grid points, we added the following radial exchange between non-wall grid points:

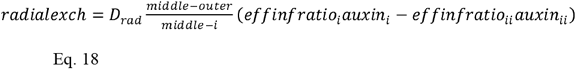

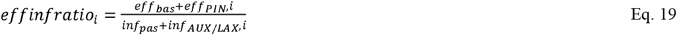

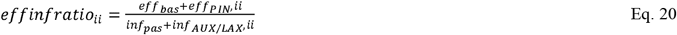

here the *effinfratio’s* reflect that the fraction of cell internal auxin available for exchange depends on the ratio between cell efflux and influx, the higher that ratio the more easily the auxin is lost to the outside where it can be exchanged with other cells. As for the passive radial fluxes, the transport rate is scaled with the inverse of the distance between the symmetrical positions.

### Staggered cell wall settings

To investigate whether the priming dynamics we observe in our root tip model critically rely on the somewhat artificial parallel cell wall positioning between neighboring cell files in our model we performed alternative simulations. Here, we incorporated the fact that for the vasculature the elongation zone starts closest to the root tip, while for the pericycle it starts furthest from the root tip, as well as differences in cell height (epidermal and endodermal cells 10, vasculature cells 12, and other cells 8 microm). Combined this results in staggered cell wall positioning between cell files.

Although we adjusted cell growth dynamics to minimize non-realistic sliding of cells past one another (see section *Cell growth, division and expansion*), sliding between neighboring cells of distinct heights can not be fully prevented in our model in which cells consist of discrete number of grid cells. As sliding effects accumulate with distance from the root tip, results from the staggered cell wall model are still quite ok close to the root tip, yet deviate more further from the root tip. Therefore, we used these alternative simulations solely to confirm that priming dynamics, occurring relatively close to the root tip, do not rely on cell wall positioning, yet refrain from using these simulations to study the PBS formation occurring further away from the root tip as we can not exclude sliding effects impacting auxin exchange between neighboring cells there.

Proper simulation of root growth dynamics, particularly in more realistic root topologies with staggered differentially sized cells between cell files, awaits the development of modeling formalisms incorporating the mechanics of symplastic root growth. While various efforts in this direction exist (Fozard et al. 2013, 2016; Marconi et al. 2021; Weise and Ten Tusscher 2019), none has yet managed to combine a realistic modeling of the particularly challenging, highly anisotropic elongation dynamics of root cells in models simultaneously describing cell-based gene expression and cell or finer-resolution based auxin dynamics to the authors knowledge.

### Kymographs

Kymographs were generated as described previously (See Figure 1B and movie 1 in (van den Berg et al. 2021)). Briefly, we draw a single grid point wide vertical line through the middle of the left pericycle cell file, plotting auxin levels along the length of this 1D line using a a color gradient to represent auxin levels. By plotting auxin levels along this 1D line every 100 seconds and concatenating these 1D lines in the vertical direction a space-time plot of auxin dynamics is generated.

### Numerical integration and run time performance

Similar to previous root tip modeling studies by us and others (van den Berg et al. 2021; Grieneisen et al. 2007; Mähönen et al. 2014) we used an alternating direction semi-implicit integration scheme for the grid-level auxin partial differential equations (Peaceman and Rachford 1955), using integration steps of 0.4s and a spatial integration step of Δx= 2μm. For the cell-level gene expression ordinary differential equations standard Euler forward integration was applied. The code of the model was written in C++, simulations were run on 24 to 36-core workstations with Intel Xeon E5-2687W processors, resulting in a typical run-time of ~12 hours for a simulation representing 6 days of plant growth.

## Acknowledgements

We thank JanKees van Amerongen for technical support and Monica Garcia Gomez for valuable comments on the manuscript.

## Competing interests

No competing interests declared.

## Funding

JTS was funded by grant nr 737.016.012, and TvdB and KtT were funded by grant nr 864.14.003 of the Netherlands Scientific Research Organization (NWO).

## Data availability

The source code necessary to perform all simulations underlying the figures in this manuscript is will be made publicly available upon publication.

## Supplementary Figure Legends

**Suppl Fig 1.**
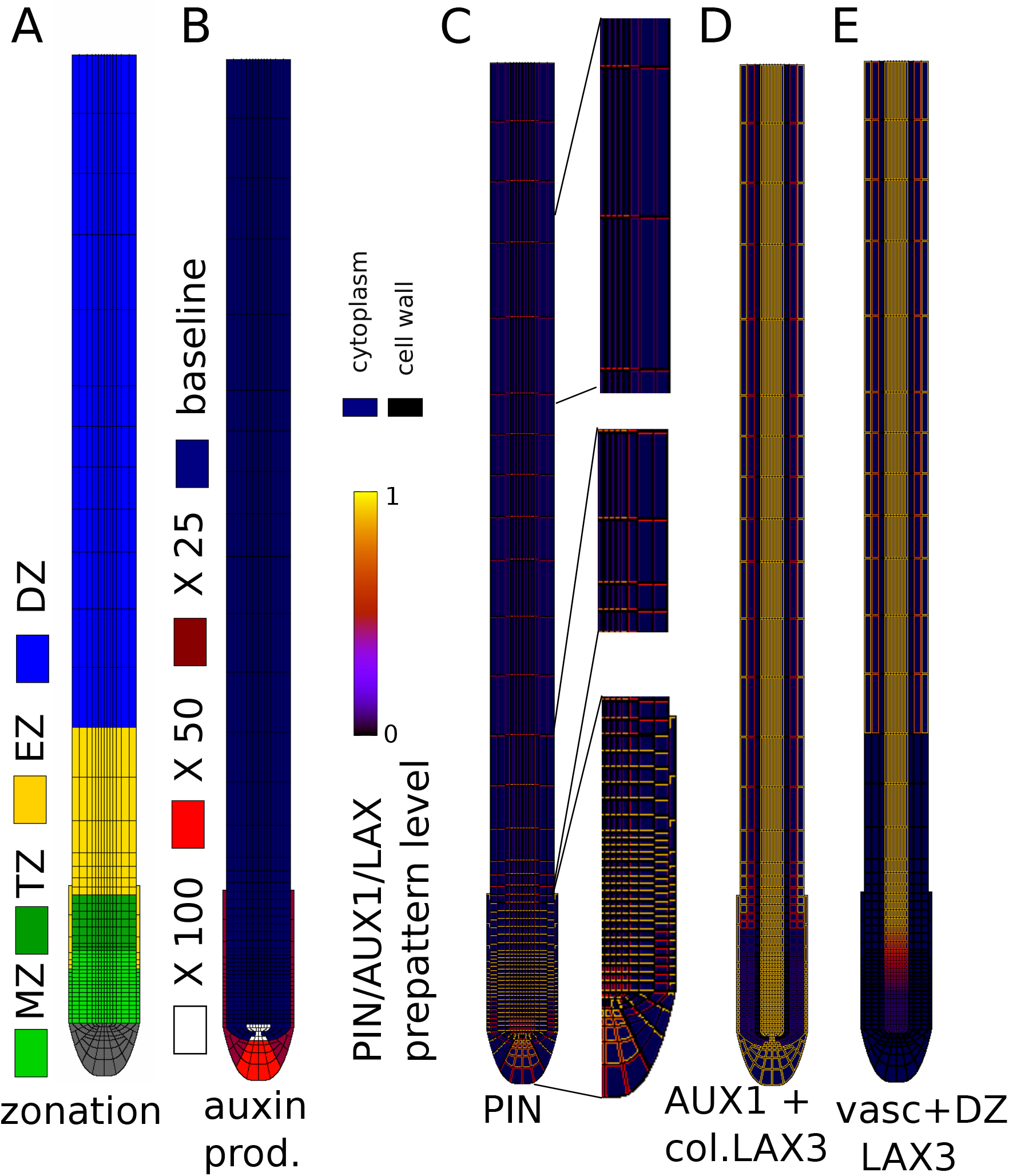
Overview of model layout. A) Root tip developmental zonation consisting of a proper meristematic zone in which cells divide (MZ), a transit amplifying zone (TZ) where cells still grow cytoplasmatically but no longer divide, an elongation zone (EZ) where cells undergo rapid vacuolar expansion and a differentiation zone (DZ) where cells undergo terminal differentiation. B) Cell type specific differences in auxin production rates given as fold increase relative to the baseline auxin production rate. C) PIN prepattern. D) AUX1 and columellar LAX3 prepattern. E) Vascular and differentiation zone epidermal and cortical LAX3 prepattern. Note that prepattern membrane levels signify maximum levels, they are multiplied by (normalized) gene expression level to determine the actual, active membrane levels.

**Suppl Fig 2.**
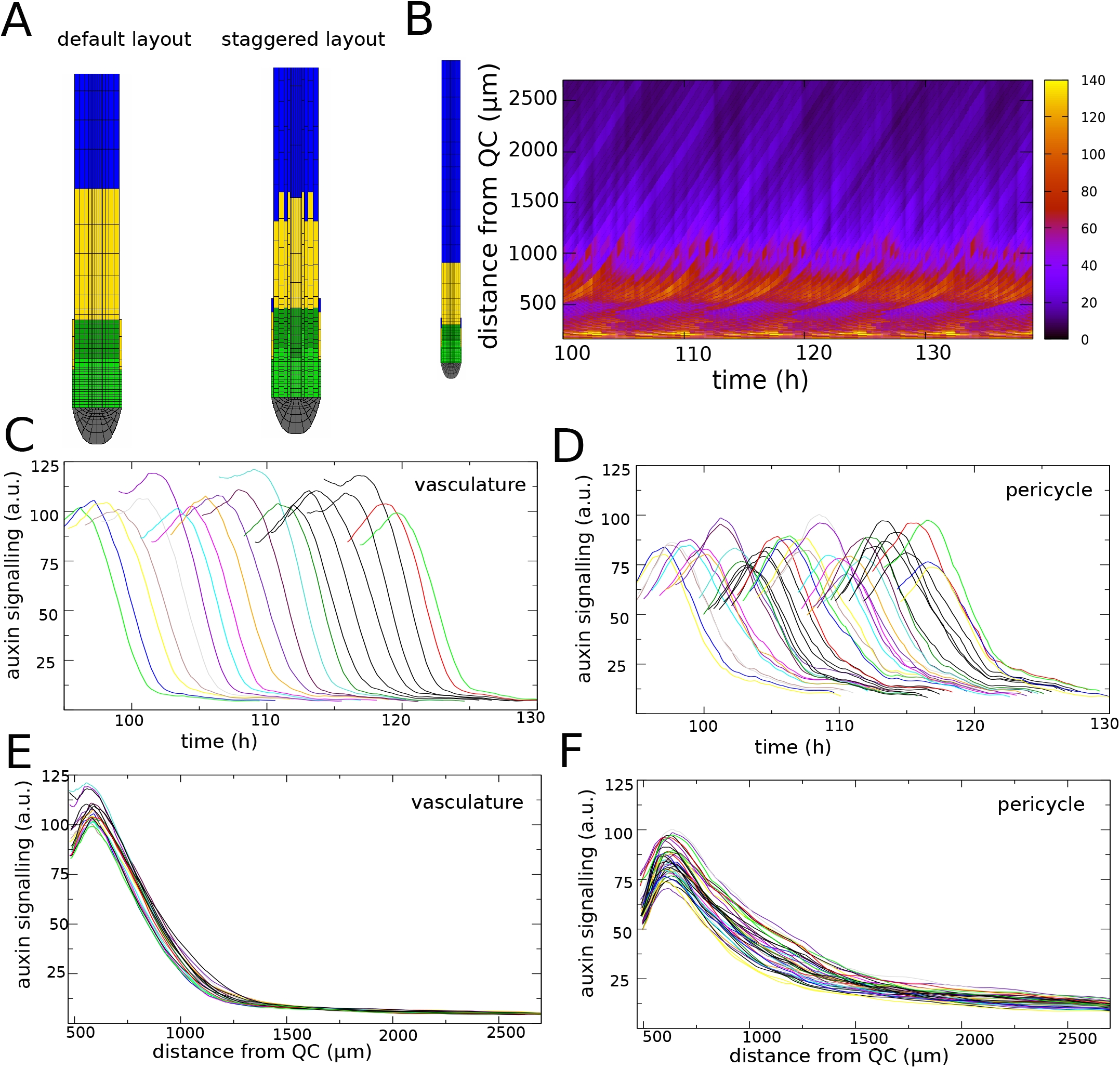
Priming under staggered cell wall positioning. A) comparison of cell layout in case of similar cell sizes and similar developmental zone boundaries resulting in parallel cell wall locations across cell files (left) and in case of different cell sizes and transition zone boundaries resulting in staggered cell wall locations across cell files (right). B) Kymograph of pericycle auxin dynamics for the staggered cell wall model. All other settings are equal to those applied in Fig 2. C,D) Vascular (C) and pericycle (D) auxin dynamics as a function of time. E,F) Vascular (E) and pericycle (F) auxin dynamics as a function of distance from the root tip. Note that due to cell size differences, within the same time window a different number of cells is traced for these two tissue types, and that larger vascular cells being next to a variable number of variably positioned pericycle cells result in more irregular priming dynamics.

**Suppl Fig 3.**
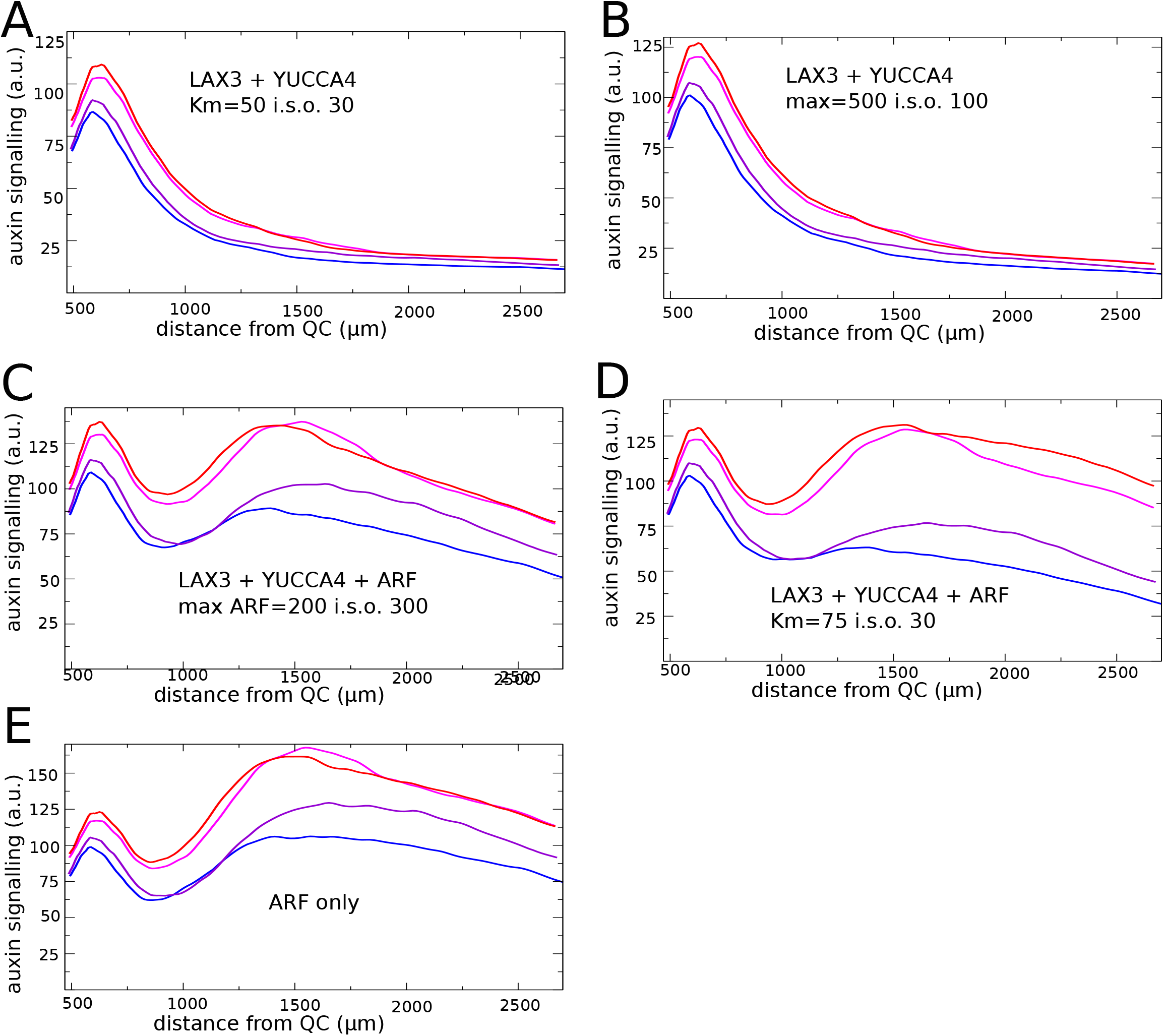
Positive feedback for alternative parameter settings. A) Pericycle auxin signalling for only LAX3 +YUCCCA4 positive feedback for Km=50 instead of Km=30 (Fig 3B), indicating limited effect of Km on effectiveness positive feedback. B) Pericycle auxin signalling for only LAX3 + YUCCA4 feedback for maximum expression 500 instead of 100 (Fig 3B), indicating limited effect of maximum expression on effectiveness positive feedback. C) Pericycle auxin signalling for LAX3+YUCCA4+ARF feedback for max ARF expression 200 instead of 300 (Fig 3C), indicating a quantitative effect but otherwise similar behavior. D) Pericycle auxin signalling for LAX3+YUCCA4+ARF feedback for Km=75 instead of 30 (Fig 3C), indicating this does not significantly enhance differentiation between cells receiving different strengths of priming signal yet reduces strength of the secondary auxin signalling response. E) Pericycle auxin signalling in absence of auxin signalling induced YUCCA4 and additional LAX3 expression.

**Suppl Fig 4.**
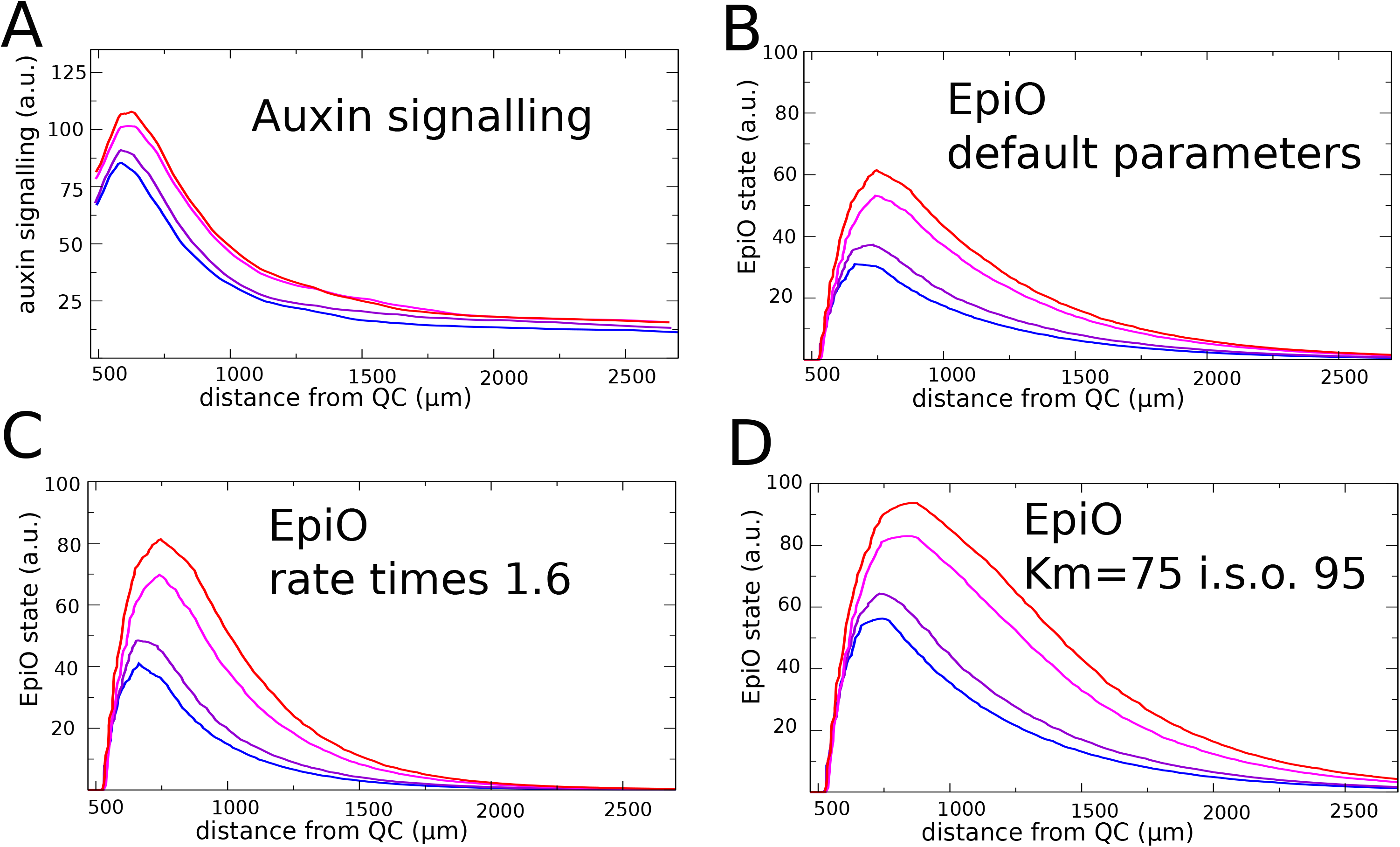
EpiO dynamics for alternative parameter settings. A) Non-normalized auxin signalling dynamics as a function of distance corresponding to the normalized auxin dynamics shown in Fig 4C. B) Non-normalized EpiO dynamics corresponding to the normalized EpiO dynamics shown in Fig 4C. C) EpiO dynamics for non-default model settings where the production and degradation rates of EpiO where increased 1.6 fold. D) EpiO dynamics for non-default model settings where the Km for AuxinSignalling induced increase of EpiO state was changed from 95 to 75.

**Suppl. Fig 5.**
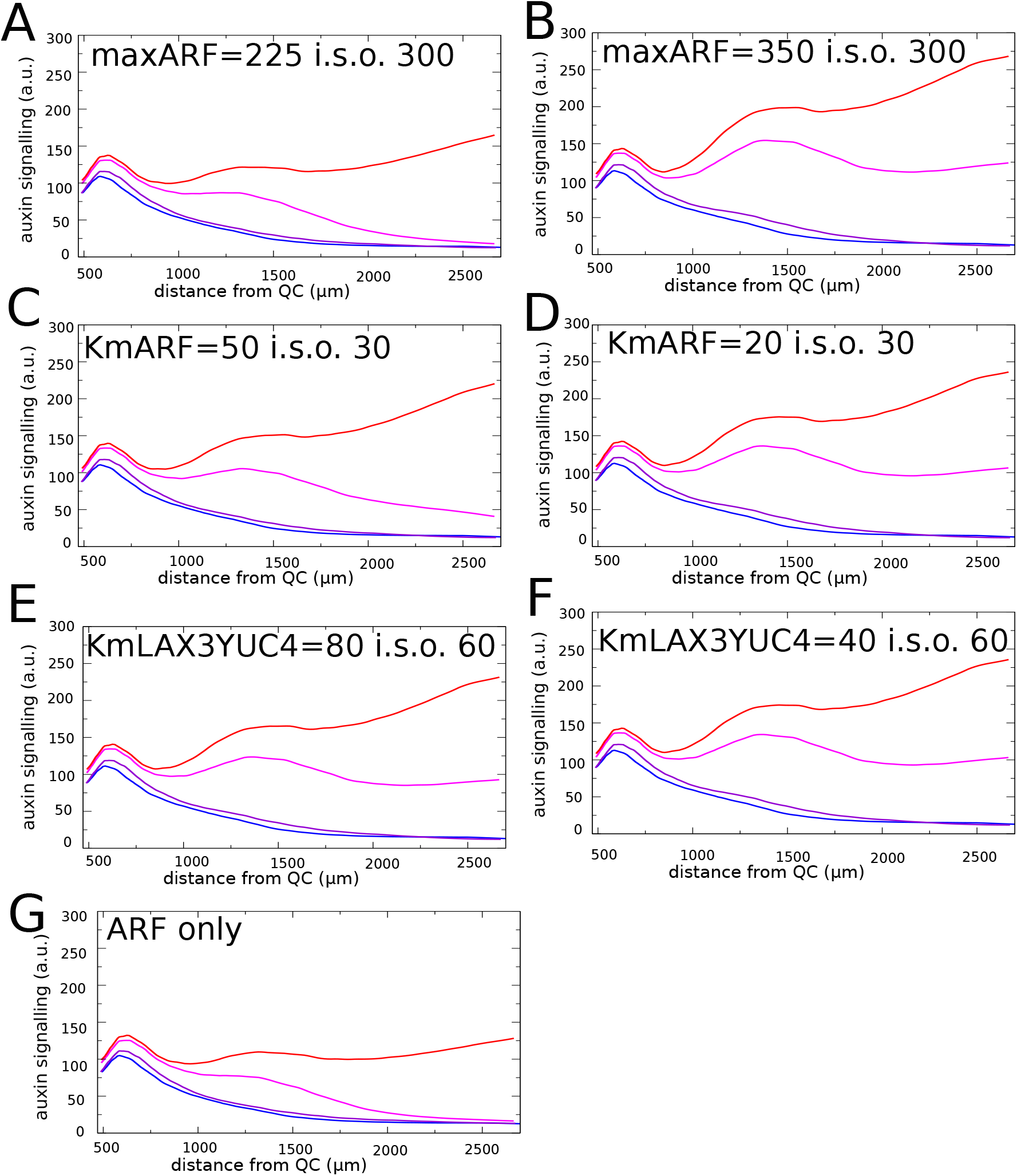
Auxin signalling dynamics in EpiO + positive feedback model for alternative parameter settings. A, B) Decreased (A) and increased (G) maximum ARF expression, reduces respectively enhances overall auxin signalling levels. In case of a decrease formation of a second stable PBS does not occur. C,D) Increased (C) and decreased (D) Km for auxin signalling induced ARF expression reduces, respectively enhances overall auxin signalling levels. In case of an increase formation of a second stable PBS does not occur. E,F) Increased (E) and decreased (F) Km for auxin signalling induced LAX3 and YUCCA4 expression hardly effects overall auxin signalling dynamics. G) Auxin signalling dynamics in case only upregulation of ARF, but not LAX3 or YUCCA4 occurs.

**Suppl Fig. 6.**
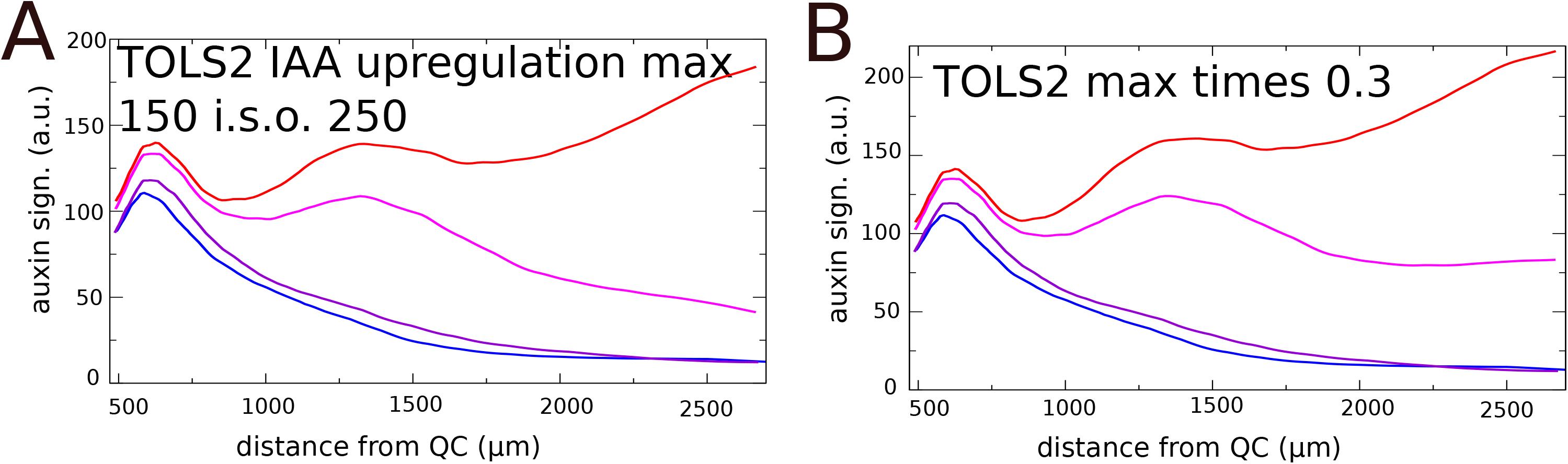
TOLS2 signalling strength affects lateral inhibition efficiency. A) Reduction of the TOLS2 signalling effect on IAA induction with 40% reduces speed of secondary PBS inhibition. B) Reduction of the TOLS2 signalling level by 66.7% abolishes secondary PBS inhibition.

**Suppl Fig. 7.**
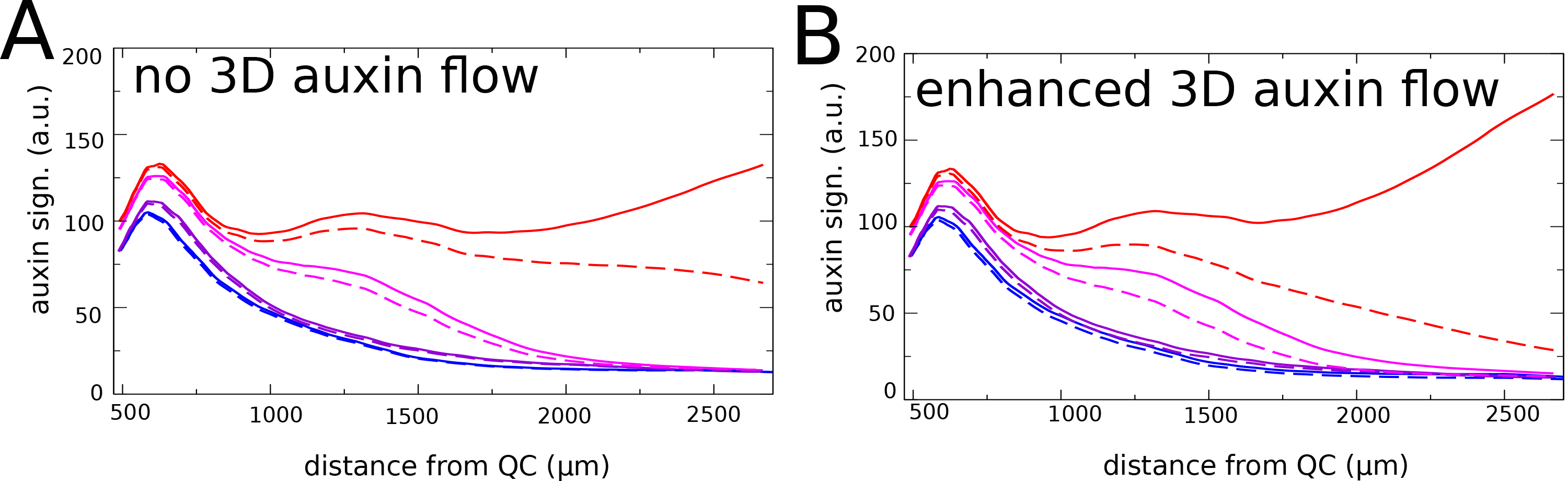
Symmetry breaking for alternative parameter settings. A) Default model settings. B) Increase of LAX3 Km from 60 to 75. C) Increase of TOLS2 mediated maximum IAA induction by 40%. D) Initial assymmetry from 10% left-right difference in LAX3.

**Suppl Fig. 8.** 3D auxin flows enhance symmetry breaking. A) Absence of 3D auxin flows slows down repression of low AUX1 side PBS. B) Enhancement of 3D auxin flows enhances repression of low AUX1 side PBS. (For details see Methods)

## Supplementary Movie Legends

**Suppl. Movie 1** Dissipation of auxin signalling in a priming only simulation. The movie corresponds to the results shown in Figure 2.

**Suppl. Movie 2** Auxin concentration, ARF, Auxin signalling, YUCCA 4 and LAX3 dynamics for the direct positive feedback model settings. The movie corresponds to the results shown in Figure 3C-D.

**Suppl. Movie 3** Auxin concentration, EpiO, ARF, Auxin signalling, YUCCA 4 and LAX3 dynamics for the time integrated positive feedback model settings. The movie corresponds to the results shown in Figure 4 A-E and Figure 5 A-B.

**Suppl. Movie 4** Auxin concentration, EpiO, ARF, Auxin signalling, YUCCA 4 and LAX3 dynamics for the time integrated positive feedback model settings incorporating TOLS2-RLK7-PUCHI mediated lateral inhibition. The movie corresponds to the results shown in Figure 6A-D.

**Suppl. Movie 5** Auxin concentration, EpiO, ARF, Auxin signalling, YUCCA 4 and LAX3 dynamics for the time integrated positive feedback model settings incorporating TOLS2-RLK7-PUCHI mediated lateral inhibition and AUX1 induced symmetry breaking. The movie corresponds to the results shown in Figure 7A-B.

## Notes

### Competing Interest Statement

The authors have declared no competing interest.

